# Spatial and spectral mapping of traffic-related air pollution (TRAP) nanoparticles in relation to plaques and inflammatory markers in an Alzheimer disease model

**DOI:** 10.1101/2025.07.01.662638

**Authors:** Hannah J O’Toole, Anchaleena James, Nathifa Nasim, Dustin J Hadley, Elizabeth J Hale, Qing He, Keith J Bein, Anthony Venezuela, Tatu Rojalin, Brittany N Dugger, Anthony S Wexler, Pamela J Lein, Randy P Carney

## Abstract

Chronic exposure to traffic-related air pollution (TRAP) is linked to increased risk of neurodegenerative diseases, including Alzheimer disease (AD). Ultrafine particulate matter (UFPM) is a suspected driver of TRAP neurotoxicity, but its spatial interactions with AD pathology remain poorly defined. We investigated the distribution, composition, and pathological context of TRAP-derived UFPM in the hippocampus of TgF344-AD rats chronically exposed to TRAP or filtered air (FA) for 14 months. Using a multimodal imaging workflow that combines enhanced darkfield hyperspectral imaging (EDF-HSI) with confocal immunofluorescence for microglia (CD68/Iba1) and amyloid beta (Aβ) plaques (Thioflavin S), we mapped the localization and spectral properties of UFPM *in situ*. UFPM accumulation was elevated in TRAP-exposed females, suggesting sex-specific vulnerability in blood-brain barrier (BBB) permeability or particle retention. Particles near plaques showed red-shifted spectral signatures, consistent with biochemical transformation. Dimension reduction revealed clustering of particle spectra by TRAP exposure and plaque proximity. However, UFPM was rarely found within plaques or microglia, implying indirect neuroimmune modulation. These findings highlight a novel spatial and spectral imaging approach for characterizing environmental nanoparticle interactions in the brain and suggest that chronic TRAP exposure may influence AD-related inflammation and pathology in a sex-and region-dependent manner.

**Synopsis:** This study shows that chronic traffic-related air pollution alters ultrafine particulate matter deposition and neuroinflammation in a rodent Alzheimer model, revealing region- and sex-specific vulnerability in the brain’s response to environmental exposure.

## Introduction

Traffic-related air pollution (TRAP) is a complex mixture of airborne particulate matter (PM) and gas-phase pollutants produced by vehicles^1,2^. Epidemiological studies link chronic exposure to TRAP with increased risk of age-related dementia, including Alzheimer disease (AD).^3,2,4^

One feature of TRAP is an elevated amount of fine and ultrafine particulate matter (UFPM, ≤100 nm), which, unlike larger particles, can more readily access the brain post-inhalation.^5^ These nanoparticulates are also less effectively cleared by alveolar macrophages and can traffic adsorbed toxic compounds into circulation, provoking inflammatory and oxidative stress responses in both the lungs and the central nervous system.^6,7^ Through such pathways, inhaled TRAP constituents may reach and impact sensitive brain regions, potentially initiating or exacerbating neurodegenerative processes.^3^

Consistent with this model, toxicological studies in animals suggest TRAP exposure can exacerbate key pathological features of AD, including oxidative stress, neuroinflammation, amyloid beta (Aβ) plaque accumulation, and tau hyperphosphorylation in the brain.^3,8^ Neuroinflammation has emerged as a potential mechanistic link between air pollution and neurodegeneration, with microglial activation proposed as a mediator of TRAP’s effects on the aging brain.^3,9^ However, most prior animal studies focused on isolated components of TRAP or employed acute, high-concentration exposures that far exceed ambient levels, limiting their relevance to human chronic exposure.^3^ These limitations underscore the need for models that more faithfully replicate real-world TRAP exposures over extended periods.

Our recent preclinical study addressed the exposure gap by collecting TRAP from a heavily trafficked vehicular tunnel and delivering it to rats housed in an adjacent vivarium unchanged and in real-time for up to 14 months.^8^ Notably, the particulate concentrations of the TRAP exposures in this paradigm (∼10 µg/m^3^)^10^ are similar to current air quality standards (9µg/m^3^),^11^ yet the TRAP-exposed rats exhibited markedly accelerated AD-like phenotypes ^8^ Chronic exposure to TRAP aggravated AD-like neuropathology. For example, TRAP exposure accelerated Aβ deposition and promoted microglial activation in male and female TgF344-AD rats relative to filtered air controls housed in the same facility.^8^ These findings aligned with observed TRAP promotion of other AD-like phenotypes, such as hyperphosphorylated tau and neuronal cell loss, as well as accelerated cognitive impairment.^8^

Interestingly, nano-sized particulate deposits were detected within the hippocampus of TRAP-exposed animals, implicating a direct accumulation of inhaled TRAP-derived PM in rodent brain tissue.^8^ These observations raised questions about the spatial relationships between deposited TRAP particles and AD pathologic phenotypes, such as Aβ plaques and microglial activation, potentially yielding insight into the mechanistic links between pollution exposure and neurodegeneration. Here, we build upon our prior foundation by investigating how chronic TRAP exposure impacts the brain of a transgenic AD rat model at a microscopic spatial level.

In this study, we employed the TgF344-AD rat, which expresses human AD-linked mutations and recapitulates age-dependent Aβ plaque deposition and neuroinflammatory responses observed in the human condition.^12,13^ TgF344-AD rats are hemizygous for two human transgenes, APPswe and PSEN1ΔE9, which are linked and inherited in a simple Mendelian fashion that manifests the full AD phenotype.^13^ In these rats, we examined the hippocampus, a vulnerable region in early AD, after prolonged exposure to either TRAP or filtered air (FA). A novel combination of enhanced darkfield hyperspectral imaging (EDF-HSI) and confocal immunofluorescence microscopy was used to map TRAP-derived PM alongside Aβ plaques and select microglial markers of neuroinflammation (**Figure 1**). EDF-HSI enabled visualization of light-scattering PM within brain sections, while immunofluorescent labeling identified CD68-positive (CD68^+^) and/or Iba1-positive (Iba-1^+^) microglia, as well as thioflavin S-positive (ThioS^+^) Aβ plaques. This integrated approach provides high-resolution spatial mapping of inhaled particles relative to microglia and plaques *in situ*, offering a unique readout of TRAP’s neurotoxic effects. By correlating the distribution of TRAP-derived particles with localized neuroinflammatory and amyloid pathology, our study aims to advance the understanding of how chronic TRAP exposure may initiate or exacerbate AD pathology and to demonstrate the utility of improved chronic exposure models and imaging methodologies in environmental neurotoxicology research.

**Figure 1.**
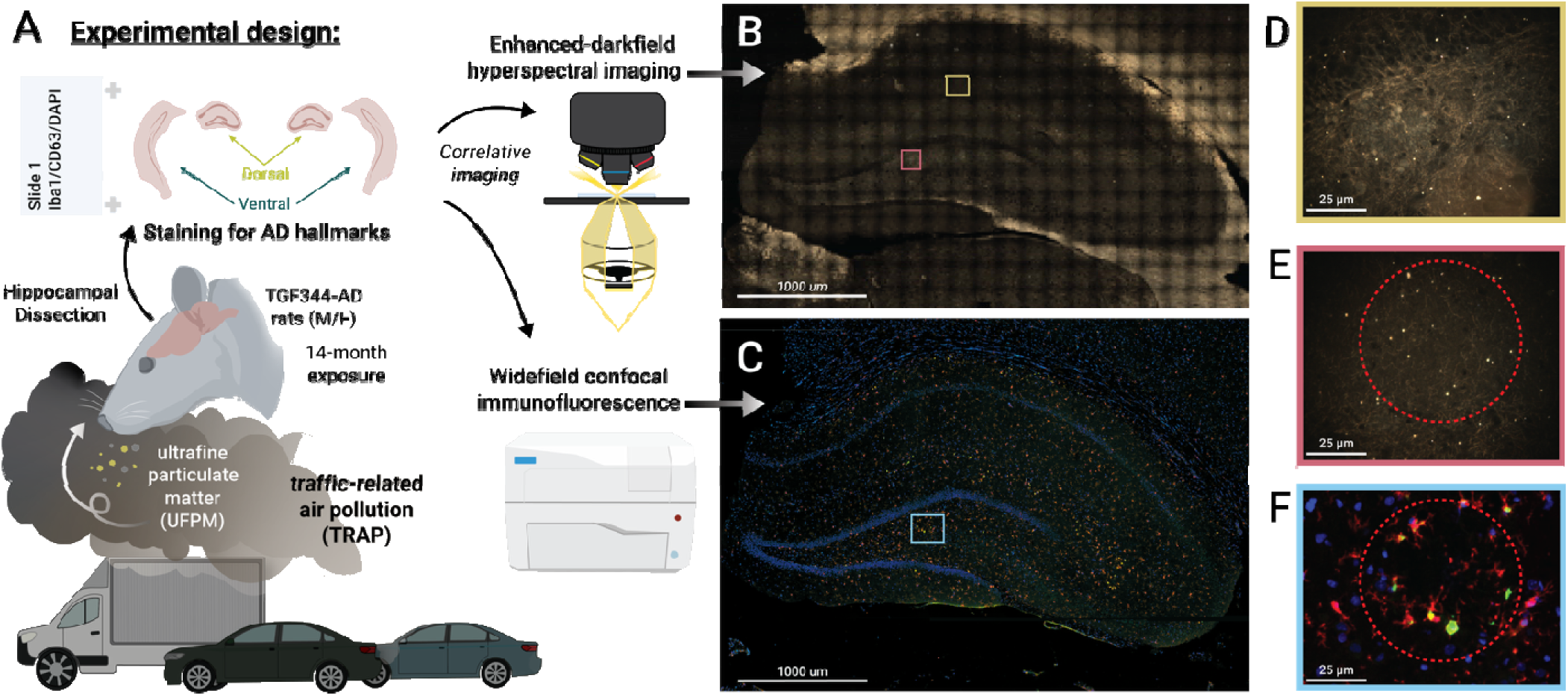
Multimodal imaging workflow for mapping TRAP-derived particulate matter and Alzheimer disease (AD) pathology in TgF344-AD rats. (A) Schematic overview of the experimental approach. TgF344-AD rats were chronically exposed to traffic-related air pollution (TRAP) or filtered air (FA) for 15 months. Brain tissue was harvested and processed for multimodal imaging: (1) Enhanced darkfield hyperspectral imaging (EDF-HSI) was used to localize and spectrally profile individual particulate matter (PM) within dorsal and ventral hippocampal sections; (2) Adjacent sections were immunostained with AlexaFluor488 for CD68 (green) and AlexaFluor568 for Iba-1 (red) to localize phagocytic microglia, stained with Thioflavin S (shown in Figs. 3 and 5) for Aβ plaques, and then imaged using high-resolution tiled confocal microscopy. (B) Representative darkfield image of a hippocampal section from a TRAP-exposed female TgF344-AD rat, showing PM distribution across the dorsal (shown) and ventral regions. Colored boxes indicate regions of interest for high-magnification views shown in (D–F). (C) Corresponding widefield confocal immunofluorescence image of a matched section stained for DAPI (blue, nuclei), CD68 (green), and Iba-1 (red). (D) Representative high magnification (40X) darkfield image illustrating the presence of PM particles (bright white puncta) in the parenchyma. (E) Example region containing a dense plaque (dotted red circle) with adjacent, red-shifted PM accumulation. (F) Overlay of confocal immunofluorescence and hyperspectral imaging from the same region as (E), highlighting the spatial proximity of PM (bright white/yellow puncta) to Iba-1^+^ microglia (red) and CD68^+^ phagocytic cells (green), with little evidence of direct particle internalization.

## Results

### TRAP exposure increases hippocampal particle burden with sex-dependent differences in accumulation patterns

To assess whether chronic TRAP exposure leads to increased PM accumulation in the brain, we performed EDF-HSI across hippocampal sections from male and female TgF344-AD rats exposed to TRAP or FA for 15 months. A total of n=24 hippocampal regions were assessed (n=12 dorsal and n=12 ventral from 6 female and 6 male brains). An anatomical visualization of our denotation of dorsal versus ventral hippocampal regions and a thorough imaging workflow is shown in **Supplemental Fig. S1**. Representative stitched EDF-HSI images from the dorsal hippocampus of two female animals (**Fig. 2A, D**) visually illustrate a significant increase in bright, scattering PM in the TRAP-exposed rat (**Fig. 2D**) relative to the FA control (**Fig. 2A**). Quantitative analysis of PM distribution (**Fig. 2B, E**) revealed that TRAP-exposed subjects showed a significantly elevated particle count (p < 0.01, student’s t-test) compared to FA controls. Qualitative examination of corresponding heatmaps in representative subjects (**Fig. 2C, F**) further confirmed this trend, with more extensive PM accumulation across the hippocampal field of view in the TRAP-exposed brain.

While these examples of the entire dorsal hippocampal region in two female rats are illustrative, downstream group-level analysis confirmed these observations. Total particle counts were quantified across 15 fields of view per hippocampal region (dorsal or ventral), and the mean count per region was used as a single data point. Each dot in **Figure 2G** represents one hippocampal region from one animal. Two-way ANOVA was performed to assess the effects of sex and exposure condition. A significant interaction between sex and exposure was observed [F(1,42) = 5.87, p = 0.0198], along with a main effect of sex [F(1,42) = 13.46, p = 0.0007]; there was not a significant main effect of exposure [F(1,42) = 3.94, p = 0.0538]. The significant interaction indicated that the effect of exposure conditions on PM counts differed by sex. Post hoc analysis (Tukey’s multiple comparisons test) revealed that TRAP-exposed females exhibited significantly greater total PM counts compared to FA controls (p=0.0202 for FA-exposed females and p = 0.0018 for FA-exposed males) and compared to TRAP-exposed males (p =0.0007) (**Fig. 2G**).

**Figure 2.**
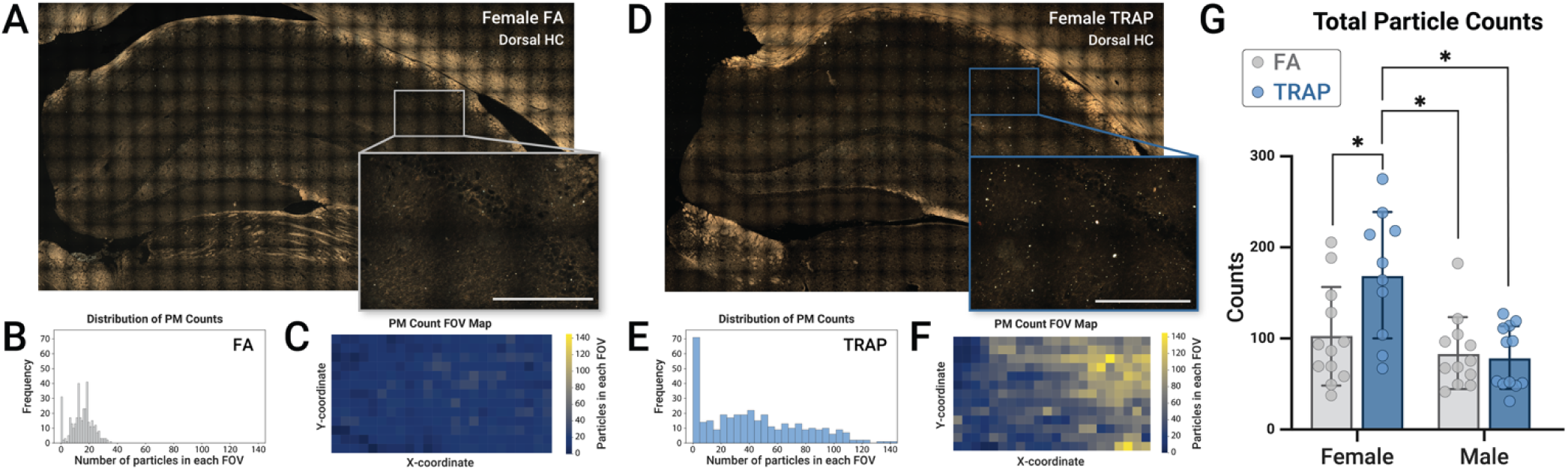
Quantification of TRAP-derived particle accumulation in the hippocampus reveals sex-dependent retention and diffuse tissue distribution. (A, D) Stitched widefield EDF-HSI imaging of the dorsal hippocampus from two FA-exposed (A) and TRAP-exposed (D) female TgF344-AD rats. Insets show magnified regions highlighting white-scattering TRAP PM (Scale bars: 100 µm). (B, E) Histograms of particle count per field of view (FOV) show distributions of particle accumulation, with more particles overall in the TRAP-exposed animal. (C, F) Heatmaps of spatial particle density reveal diffuse but elevated local concentrations of PM in the TRAP-exposed animal. (G) Total PM counts pooled across dorsal and ventral hippocampal regions (area normalized) were quantified from 15 FOVs per region and averaged to yield one value per hippocampal region (represented by a single dot from one animal). TRAP-exposed females exhibited significantly higher particle burden than all other groups (*p < 0.05, Two-way ANOVA with post hoc Tukey’s). Bars show mean ± SD.

### TRAP exposure does not exacerbate A**β** plaque burden but does reveal regional and sex-dependent vulnerability

To assess the impact of chronic TRAP exposure on amyloid pathology, we quantified ThioS^+^ plaques in the dorsal and ventral hippocampal regions of TgF344-AD rats. ThioS^+^ regions from confocal imaging were quantified using ImageJ thresholding and normalized to the hippocampal area. Representative ThioS-stained dorsal hippocampal regions from two female rats (**Fig. 3A, B**) visually illustrate the presence of Aβ plaque burden across both FA and TRAP exposures. The full-resolution imaging dataset is included as **Supplementary** Figure 2, which displays representative widefield ThioS-stained sections from both dorsal and ventral hippocampus regions, matched by exposure and sex. Such images were used to generate the plaque quantification results presented in **Figure 3C-E**. Plaque counts were quantified in 30 distinct hippocampal regions from n = 24 animals; each marker in **Figure 3C-E** represents one analyzed hippocampal region (dorsal or ventral) from one animal. Aβ plaque burden was evaluated across groups by comparing area-normalized plaque counts to sex, hippocampal region (dorsal vs. ventral) and exposure condition (FA vs. TRAP) using a two-way ANOVA (**Fig. 3C**). A significant main effect of sex/hippocampal region was observed [F(3,22) = 40.42, p < 0.0001], accounting for 79.17% of the total variance. However, there was no significant effect of total plaque burden between rats exposed to FA and those exposed to TRAP for 14 months [F(1,22) = 0.14, p = 0.7099] (**Fig. 3C**). No interaction between factors was observed. Post hoc Tukey’s multiple comparisons revealed several significant differences (adjusted p < 0.05). Plaque burden was consistently higher in the dorsal versus ventral hippocampus across both sexes and exposure conditions (p < 0.0001 for most comparisons). Male dorsal regions exhibited significantly higher plaque burden than all other male and female groups across both conditions. Given the lack of observed significant effect of TRAP exposure on Aβ plaque burden in the full model (**Fig. 3C**), we next collapsed across the exposure conditions to isolate the effects of hippocampal region (**Fig. 3D**) and sex (**Fig. 3E**).

To assess Aβ plaque burden across hippocampal subregions independent of sex, a two-way ANOVA was conducted with region and exposure as factors (**Fig. 3D**). A significant main effect of hippocampal region was observed [F(1,26) = 85.23, p < 0.0001], with dorsal hippocampal regions showing consistently greater plaque burden than ventral regions, confirmed with post hoc Tukey’s multiple comparisons.

We then examined sex differences in Aβ plaque distribution independent of exposure condition by comparing area-normalized plaque counts with sex and hippocampal region as factors (**Fig. 3E**). A significant main effect of the hippocampal region was observed [F(1,26) = 108.9, p < 0.001], with higher plaque burden in dorsal regions. Post hoc comparisons revealed significantly higher plaque counts in the dorsal versus the ventral hippocampus overall across sexes (p < 0.0001). Male dorsal hippocampal regions showed significantly greater plaque burden than female dorsal (p = 0.0392), but no sex difference was detected in ventral regions (p = 0.1512) (**Fig. 3E**). These findings suggest chronic TRAP exposure does not significantly exacerbate Aβ plaque accumulation in this model. Instead, the data reveal an intrinsic regional neuroanatomical and sex-dependent pattern of vulnerability, with more dorsal plaque burden in males, regardless of exposure.

**Figure 3.**
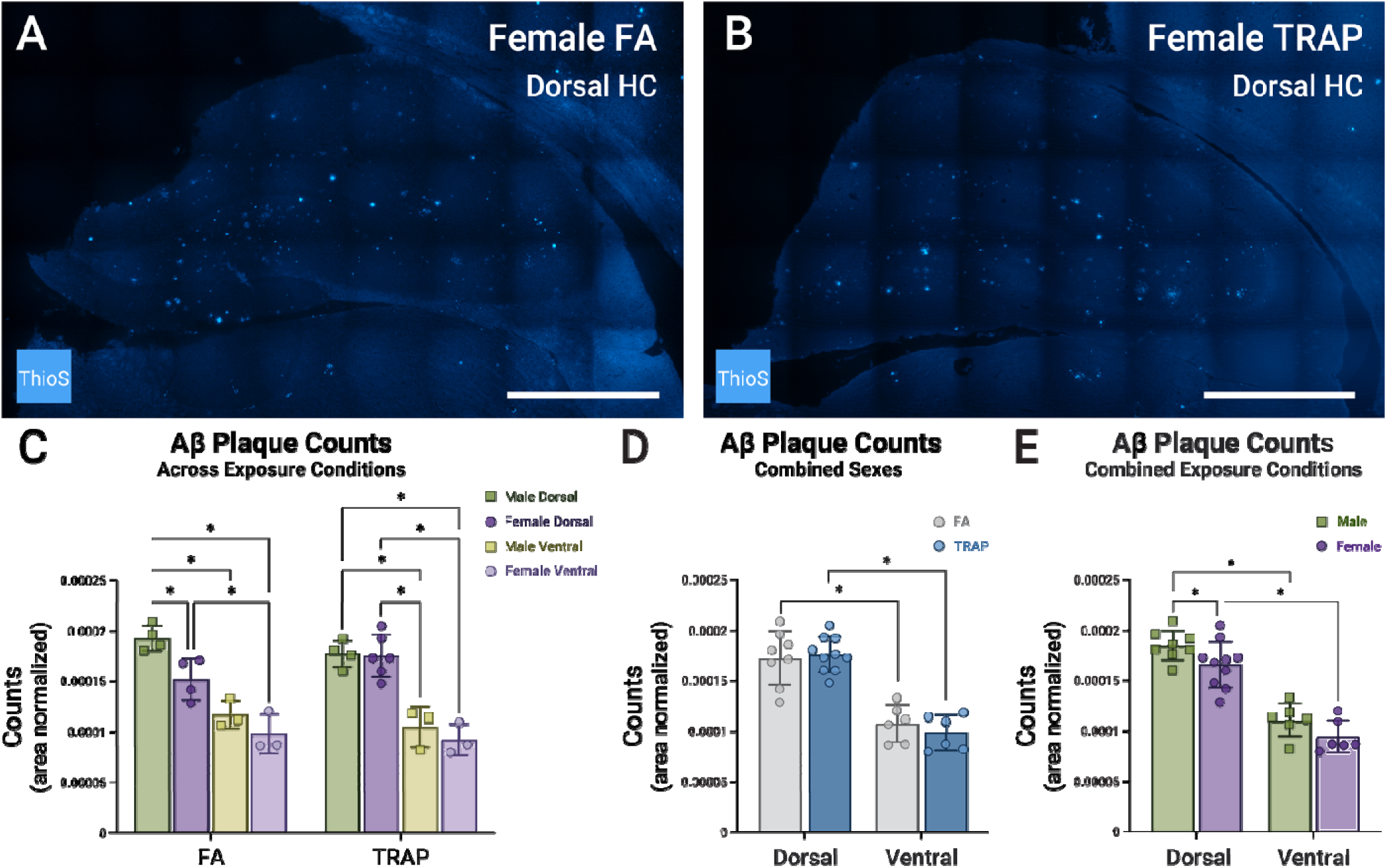
Aβ plaque burden in TgF344-AD rats shows regional and sex-dependent differences, but no significant effect of TRAP exposure. (A–B) Representative Thioflavin S-stained dorsal hippocampal sections from female TgF344-AD rats after 14 months of (A) FA or (B) TRAP exposure. Fluorescent Aβ plaques are readily visible throughout the hippocampus, with higher density in the dorsal region compared to the ventral (normalized to respective areas in pixels^2^). (C) Area-normalized Aβ plaque counts were stratified by sex, exposure condition, and hippocampal subregion. Each point represents one hippocampal region (dorsal or ventral) from one animal. A two-way ANOVA revealed a significant main effect of sex/region [F(3,22 = 40.42, p < 0.0001], but no effect of exposure [F(1,22) = 0.14, p = 0.7099] or interaction. Post hoc Tukey’s multiple comparisons showed greater Aβ plaque burden in the dorsal vs. ventral regions, and significantly higher burd n in male (green squares) dorsal hippocampus compared to all other groups (* p< 0.05). (D) When data were collapsed across sex, there was a significant main effect of hippocampal region [F(1,26 = 85.23, p < 0.0001], but no effect of exposure or interaction. Dorsal plaque burden was significantly higher in both FA (grey) and TRAP (blue) exposed animals (*p< 0.05, 2-way ANOVA, post hoc Tukey’s). (E) Collapsed across exposure conditions, there were significant main effects of hippocampal region [F(1,26) = 108.9, p < 0.0001] and sex [F(1,26) = 0.6.36, p = 0.0182], but no interaction was observed. Post hoc Tukey’s multiple comparisons test showed that the dorsal hippocampus retained a significantly higher plaque burden than the ventral hippocampus for both sexes, with males (green squares) exhibiting more plaques than females (purple circles) in the dorsal hippocampus (*p < 0.05); no sex difference was observed in ventral regions. Collectively, these data show that Aβ plaque deposition in TgF344-AD rats is more pronounced in the dorsal hippocampus and males, but is not significantly altered by chronic TRAP exposure. Bars show mean ± SD. Scale bars: 1000 μm.

### Region- and sex-specific enrichment of CD68^+^ phagocytic microglia in the dorsal hippocampus

To evaluate the impact of chronic TRAP exposure on hippocampal microglial reactivity, we quantified cells co-labeled for Iba1 and CD68, biomarkers of microglial cells and phagocytosis, respectively, in dorsal and ventral hippocampal subregions of TgF344-AD rats stratified by sex and exposure condition. CD68^+^ immunoreactivity was normalized to the selected hippocampal area for each respective section. 30 hippocampal regions from 24 animals were analyzed, and each data point represents the mean CD68^+^ count from 15 fields of view per hippocampal region. A significant main effect of sex and hippocampal region [F(3,22) = 4.06, p = 0.0195], accounting for 33.6% of the total variance. However, there was no main effect of TRAP exposure [F(1,22) = 0.62, p = 0.4394], or interaction between factors [F(3,22) = 0.145, p = 0.932] (Two-way ANOVA) (**Fig. 4C**). Although CD68^+^ counts did not differ significantly between TRAP- and FA-exposed rats when analyzed across all groups (**Fig. 4C**), the significant main effect of sex and hippocampal region was observed, prompting us to collapse across sex (**Fig. 4D**) and exposure (**Fig. 4E**) to clarify the contribution of these factors with increased statistical power.

We collapsed across sex to examine the effects of exposure and hippocampal region, revealing a significant main effect of hippocampal region [F(1,26) = 10.51, p = 0.0032], with dorsal regions exhibiting higher CD68^+^ counts than ventral, but no main effect of exposure condition [F(1,26) = 0.82, p = 0.374], or interaction [F(1,26) = 0.0001, p = 0.991]. We saw a significant difference in CD68^+^ counts in the dorsal hippocampus of TRAP-exposed animals compared to the ventral hippocampus of FA-exposed (p = 0.028), but no other group comparisons reached significance.

We then collapsed across exposure groups to evaluate the effects of sex and hippocampal region, again revealing a significant main effect of hippocampal region [F(1,26) = 8.06, p = 0.0087], where the dorsal hippocampus consistently exhibited greater CD68^+^ microglial cell density than the ventral hippocampus. Post hoc comparisons confirmed dorsal enrichment of CD68^+^ counts in females (p = 0.0328, Tukey’s), with no other comparisons reaching significance (**Fig. 4E**).

**Figure 4.**
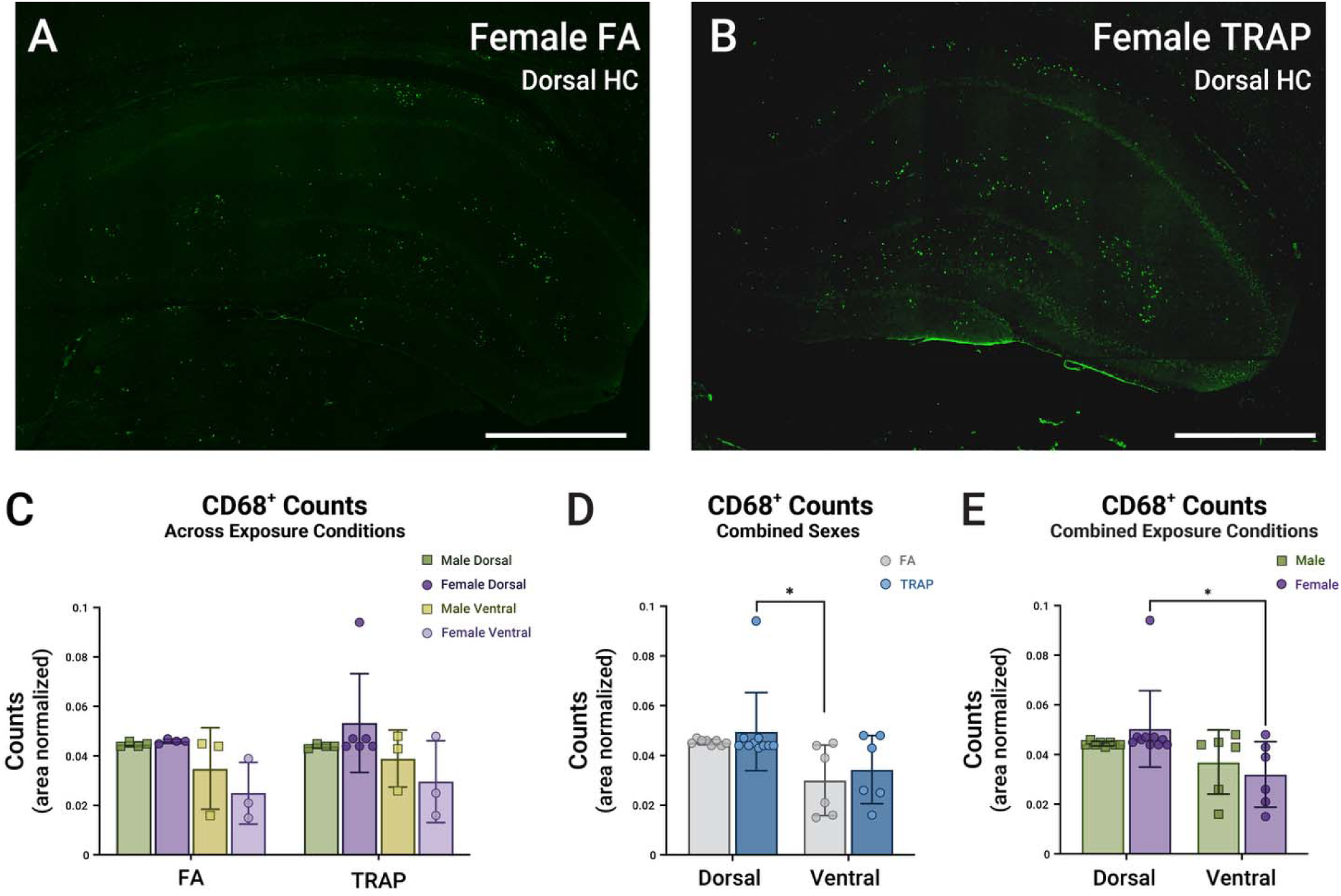
CD68^+^ microglial burden in TgF344-AD rats shows hippocampal subregion differences, but no consistent effect of TRAP exposure. (A–B) Representative CD68-stained dorsal hippocampal sections from female TgF344-AD rats after 14 months of (A) FA or (B) TRAP exposure. (C) Quantification of cells co-labeled for CD68 and Iba1 in dorsal and ventral hippocampus (normalized to respective areas in pixels^2^), stratified by sex, hippocampal region, and exposure condition. Two-way ANOVA revealed a significant main effect of sex/region F[(3,22) = 4.06, p = 0.0195], but no main effect of exposure or interaction. While no significant differences were observed across exposure groups overall, TRAP-exposed females showed a modest elevation in dorsal hippocampal CD68^+^ signal compared to other subgroups, with one notable outlier. (D) When collapsed across sex, the dorsal hippocampus showed significantly greater CD68^+^ microglial counts than the ventral region [F(1,26) = 10.51, p = 0.0032]. Post hoc Tukey’s multiple comparisons revealed significantly more CD68^+^ cells in dorsal hippocampus regions of TRAP-exposed animals than ventral regions of FA-exposed controls (*p < 0.05). (E) When collapsed across exposure condition, dorsal hippocampus regions exhibited significantly more CD68^+^ cells than ventral regions [F(1,26) = 8.06, p = 0.0087], with post hoc Tukey’s multiple comparisons testing confirming a significant difference in females (*p < 0.05). Together, these data suggest a neuroanatomic-specific neuroimmune pattern, with elevated phagocytic microglial populations in the dorsal hippocampus and sex-specific variation in regional microglial distribution in response to TRAP exposure. Bars show mean ± SD. Scale bars: 1000 μm.

### TRAP-derived particles are not enriched within AD pathological regions and remain excluded from plaques and phagocytic microglia

To assess whether PM preferentially accumulates at sites of plaques, we quantified particle abundance in hippocampal regions in ThioS^-^ areas (non-plaque regions), versus those enriched in CD68^+^/Iba1^+^ phagocytic microglia proximal to ThioS^+^ plaques. These areas were identified via confocal fluorescence microscopy, and particle counts were acquired from EDF-HSI imaging at those areas (**Fig. S2A**). Counts were collapsed across sex and hippocampal subregion (dorsal vs. ventral). A two-way ANOVA revealed, unexpectedly, no significant main effects of difference in particle counts between pathological zones and nearby plaque- and microglia-poor regions [F(1,42) = 2.28, p = 0.139]. This held for both TRAP- and FA-exposed animals [F(1,42) = 2.14, p = 0.151], suggesting that proximity to hallmark AD pathology does not predict greater PM retention, nor the reverse (**Fig. 5A**). Although TRAP-exposed animals tended to show elevated particle counts relative to FA, and plaque associated regions (ThioS^+^, CD68^+^/Iba1^+^) exhibited higher counts than plaque free regions, these trends did not reach statistical significance. Further examples of plaque and non-plaque FOVs are highlighted in **Supplemental Fig. S3**.

Moreover, spatial overlays of hyperspectral imaging and immunofluorescence revealed that particles were consistently adjacent to, but not within, either Aβ plaques or microglial cell bodies (**Fig. 5B**). Even in densely plaque-laden hippocampal subregions, particles were excluded from plaque cores and not visibly sequestered within CD68v or CD68^+^/Iba1^+^ microglia. This lack of direct incorporation was consistent across all exposure conditions and hippocampal regions.

**Figure 5.**
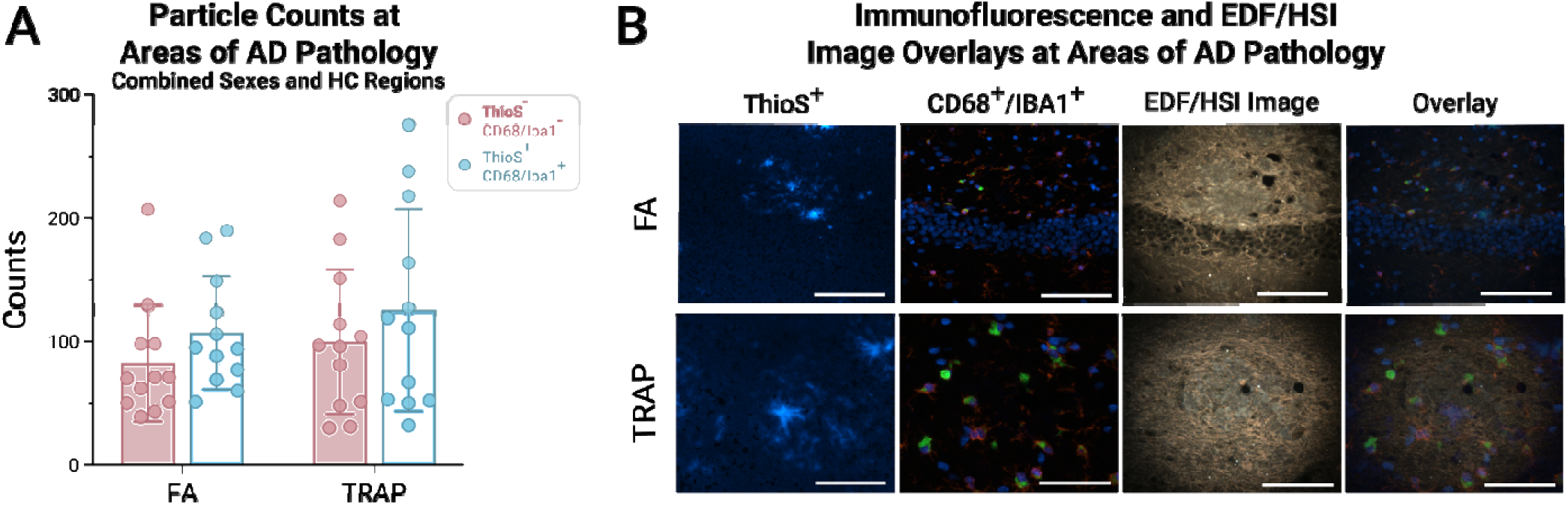
PM particles are not enriched in regions containing Aβ plaques or phagocytic microglia. (A) Quantification of PM particle counts in hippocampal regions with high AD pathological burden (defined by co-occurrence of ThioS^+^ plaques and CD68^+^/Iba1^+^ microglia, *blue bars*) versus adjacent pathology-poor areas lacking plaques or phagocytic microglia (*pink bars*). Counts were summed across 7-8 fields of view per hippocampal region and normalized per section. Each point represents one quantified hippocampal region at the individual animal level. Two-way ANOVA showed no significant main effects on particle abundance due to exposure group (TRAP vs. FA) [F(1,42) = 2.14, p = 0.151], or proximity to AD pathology [F(1,42) = 2.28, p = 0.139]. (B) Representative images from FA-exposed (*top row*) and TRAP-exposed (*bottom row*) animals show ThioS-stained plaques (*left*), CD68^+^/Iba1^+^ microglial clusters (*middle left*), and corresponding hyperspectral imaging of PM particles (*middle right*). An overlay of these FOVs (*right*) shows that despite their spatial proximity, PM particles were consistently excluded from plaque cores and were not internalized by phagocytic microglia. Bars show mean ± SD. Scale bars: 100 μm.

### Hyperspectral signatures of TRAP-derived particles reveal biochemical divergence by brain region and proximity to AD pathology

To assess whether TRAP-derived particles undergo region- or pathology-specific transformations once deposited in the brain, we performed EDF-HSI and analysis on individual particles detected within the dorsal and ventral hippocampus. **Figure 6** summarizes the spectral and dimensionality-reduced analyses across animals, stratified by exposure condition, brain region, and near or far from plaques.

Representative EDF-HSI images in TRAP-exposed rats revealed dense fields of PM distributed throughout the hippocampus (**Fig. 6A, B**). Spectra acquired from individual particles within these fields (**Fig. 6C**) showed characteristic scattering peaks near 685 nm, with TRAP-exposed particles exhibiting consistently higher intensity and broader spectral profiles than particles from FA controls (**Fig. 6D**). Averaged spectra from TRAP animals displayed a prominent red shift, particularly in the 670–700 nm region, suggestive of optical or biochemical transformations within brain tissue.

To probe underlying spectral patterns and their associations with exposure, region, and pathology, we applied dimension reduction techniques to the full spectral dataset. Principal component analysis (PCA) revealed clear separation along PC1 and PC2 between TRAP and FA particles (**Fig. 6E**), accounting for over 89% of total variance. PC loadings (**Fig. 6F**) showed the dominant contributors to spectral variance were in the ∼470 nm, ∼685 nm, and ∼870 nm ranges, with the latter peaks aligning with the observed red-shifted scattering.

We next applied uniform manifold approximation and projection (UMAP), a nonlinear method that preserves local spectral similarities, to visualize finer-grained clustering.^14^ Each dot in the UMAP plots represents an individual PM particle; particles with similar spectra are placed closer together in the two-dimensional projection. UMAP projections colored by hippocampal region (**Fig. 6G**) revealed partial clustering by dorsal vs. ventral location, suggesting region-dependent spectral differences. Supporting this, wavelength correlations with UMAP axes (**Fig. 6H**) also pointed to 685–700 nm as a dominant differentiating feature.

Particles in the ventral hippocampus exhibited greater spectral intensity and broader, red-shifted features than those in the dorsal hippocampus when stratified by anatomical region (**Fig. 6I**). These differences are not likely due to particle concentration alone, as dorsal hippocampal regions showed equal or higher particle density in other analyses.

To determine whether proximity to plaques contributes to spectral transformation, we stratified particles by their spatial distance to ThioS^+^ Aβ plaques. Particles located near plaques displayed subtle but consistent red shifts and increased intensity compared to particles further away (**Fig. 6J**), even when controlling for regional differences. Together, these findings suggest that TRAP PM not only accumulates in TgF344-AD rats but may undergo regionally distinct biochemical transformations upon interaction with pathological tissue microenvironments. These spectral signatures may serve as optical fingerprints of disease-particle interaction and could provide a platform for assessing regional vulnerability or mechanistic entry routes of airborne neurotoxicants.

**Figure 6.**
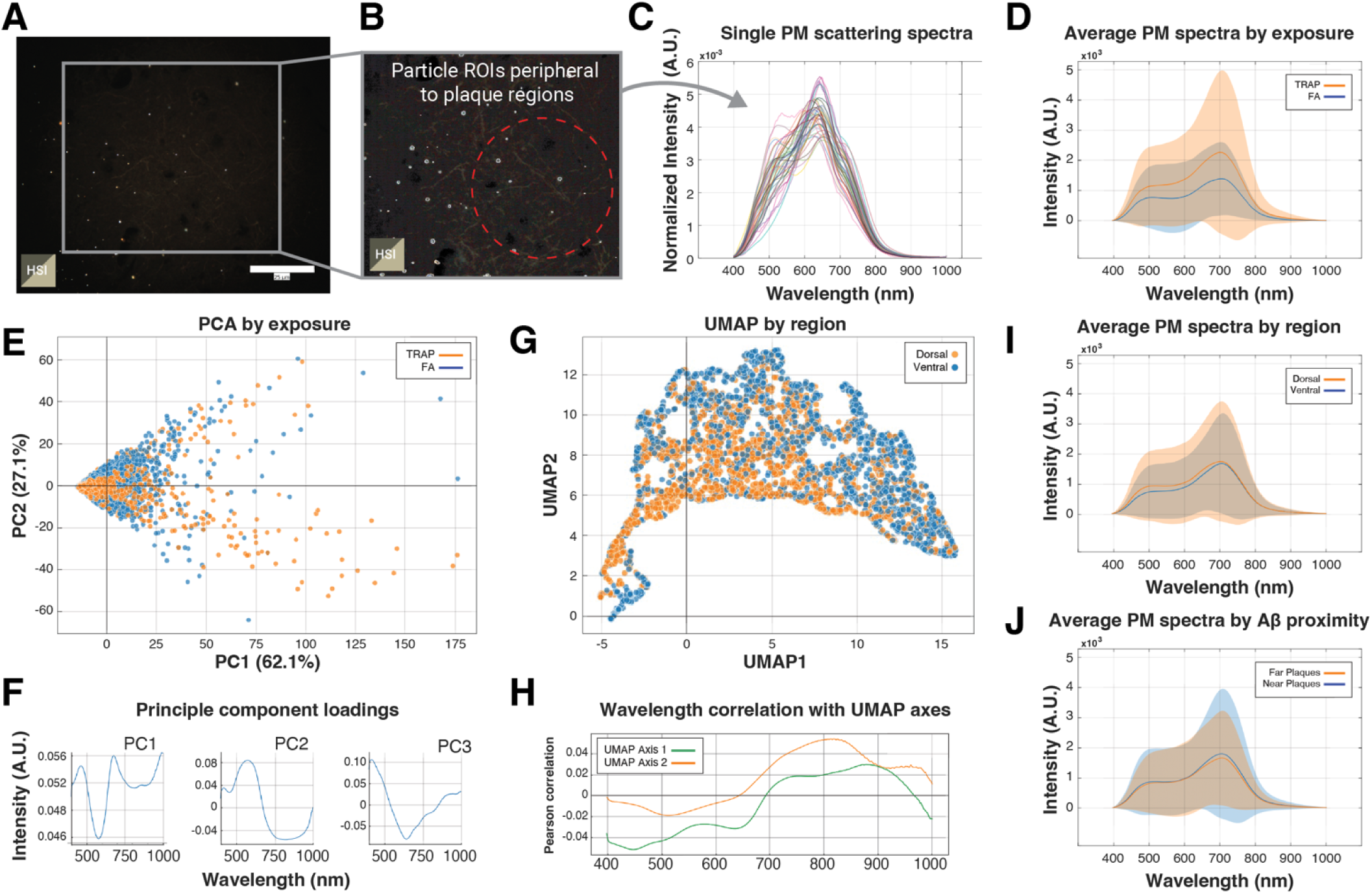
Hyperspectral analysis reveals region- and pathology-associated spectral divergence of TRAP-derived PM in the hippocampus. (A–B) Representative EDF-HSI showing PM distribution in the hippocampus. Particles, which appear here peripherally located near a plaque-dense region (outlined in red), are individually detected. (C) Example scattering spectra from individual particles. (D) Averaged spectra across all detected particles stratified by exposure condition revealed an elevated scattering intensity and red-shifted spectral peak in TRAP-exposed rats. (E) Principal component analysis (PCA) of particle spectra showed partial separation between TRAP and FA groups. (F) PCA loadings for the top three components indicated that the dominant spectral variance arose around 470, 685, and 870 nm. (G) UMAP projection colored by hippocampal region showed partial grouping of particle spectra by anatomical location. (H) Correlation between UMAP axes and wavelength further implicated the 685–700 nm region in regional particle transformation, with a red shift in ventrally-sequestered PM. (I) Direct spectral comparison by region revealed greater scattering intensity in ventral particles. (J) Many particles located near plaques showed red-shifted spectra relative to those further away, suggesting biochemical alteration near AD pathology.

## Discussion

In this study, chronic exposure to TRAP elicited significant changes in the hippocampus of TgF344-AD rats without a corresponding increase in plaques. Notably, TRAP exposure was associated with elevated CD68 expression in microglia in the dorsal hippocampus. Although traditionally interpreted as a marker of microglial activation, increased CD68 immunoreactivity in TRAP-exposed animals is now more accurately described as indicative of elevated phagocytic activity. This effect was most evident in female rats, indicating a sex-dependent neuroimmune response to TRAP. These findings are consistent with reports that particulate exposure leads to region-specific changes in microglial state, including increased phagocytic activity, in the hippocampus.^15^ Moreover, female rodents may mount a stronger central inflammatory response to airborne pollutants,^16^ aligning with epidemiological data suggesting older women are especially vulnerable to PM_2.5_-associated cognitive decline.^17^ Importantly, we observed no significant difference in Aβ plaque burden between TRAP and FA groups. This dissociation suggests TRAP’s impact on neuroinflammation in these adult TgF344-AD rats did not measurably accelerate plaque deposition over the exposure period, in agreement with prior studies where prolonged PM exposure increased neuroinflammatory markers or tau pathology without exacerbating Aβ load.^15^

A second key finding was the altered distribution and composition of PM in the brain. Female TRAP-exposed rats accumulated more brain PM than their male counterparts within the hippocampus. TRAP-exposed females had a higher density of PM in hippocampal tissue sections compared to females breathing FA for over a year (14 months), whereas male rats showed minimal difference by exposure. Although no overall sex main effect was reported in our previous analysis,^8^ our spatial mapping suggests female hippocampi may sequester more particles in proximity to plaques.

Intriguingly, our hyperspectral imaging analysis revealed that the optical signatures of particles within the hippocampus varied by brain region and their proximity to select pathological features. In particular, particles in the dorsal hippocampus, where microglial phagocytic activity (CD68 immunoreactivity) was elevated, displayed distinct optical signatures compared to those in the ventral hippocampus, suggesting differential particle processing or transformation (e.g., corona) across hippocampal subregions, and/or different particle properties (morphology, composition). Similarly, particles found adjacent to Aβ plaques or within dense microglial clusters had shifted wavelength reflectance patterns relative to particles in plaque-free regions. These observations suggest that once TRAP-derived particles penetrate the rodent brain, they may undergo physicochemical transformations influenced by the local tissue environment. Yet we observed a spatial decoupling between PM and plaques/microglia, suggesting TRAP particles may exert local effects or transformations without direct incorporation into disease structures. The lack of phagocytic internalization, even by CD68^+^/Iba-1^+^ microglia, suggests a non-canonical interaction may evade classical immune clearance pathways. Moreover, the consistent exclusion of particles from plaque cores indicates they do not passively accumulate in insoluble plaque regions. Instead, their spatial patterning supports a model in which PM interacts with the surrounding microenvironment without becoming physically sequestered.

Together, these findings argue against a model of PM acting as a passive cargo within plaques. Rather, the data support the hypothesis that TRAP PM contributes to regional vulnerability through extracellular mechanisms, possibly amplifying neuroinflammatory signaling or exacerbating BBB dysfunction in tissue already primed by disease processes.

In particular, the presence of plaques with subsequent recruitment of microglia may alter particle composition or coating: for example, a particle lodged near an Aβ plaque may adsorb plaque-associated proteins or lipids, forming a corona that changes its light-scattering properties.^18^ It is well-known that nanoparticles rapidly acquire protein coronas in biological fluids, which can modify their physicochemical behavior.^18^ Such interactions could explain the higher refractive index measured in particles from TRAP-exposed brains relative to controls.^8^ Notably, we found that TRAP-derived particles in hippocampal tissue exhibited greater maximal scattering intensity (an indicator of refractive index) than particles in FA-exposed brains. This shift could reflect oxidative modification of particle surfaces or the accretion of dense protein/lipid layers *in vivo*. In essence, the “spectral fingerprint” of brain-retained particulates appears to carry a memory of the environment they reside in, with regional differences (dorsal vs. ventral hippocampus) and pathological context (near plaque vs. far) influencing their optical characteristics. These nuanced results highlight that the impact of pollution particles in the rodent brain is not static, and once deposited, particles may be chemically altered by biological processes, potentially affecting their toxicity and persistence in the brain.

### Regional and exposure-dependent differences in plaque burden, microglial phagocytic state, and particle spectral signatures

We observed a consistent regional bias in Aβ plaque accumulation across both sexes, with the dorsal hippocampus exhibiting significantly greater plaque (ThioS^+^) burden than ventral regions, irrespective of TRAP exposure. This suggests an intrinsic dorsal vulnerability to Aβ aggregation in the TgF344-AD model, independent of environmental exposure. In parallel, CD68^+^/Iba-1^+^ phagocytic microglia were more abundant in the dorsal hippocampus, particularly in TRAP-exposed females, pointing to a sex-specific neuroimmune response that is spatially aligned with the region of higher plaque burden.

While a previous study from our group reported increased dorsal hippocampal plaque burden and CD68^+^ microglia load in TRAP-exposed TgF344-AD rats,^8^ we did not observe a significant difference in either plaque count nor CD68^+^ microglia between TRAP and FA groups in the current cohort. A few key differences between the two studies may account for this discrepancy. First, the animals used here represent a distinct exposure cohort, housed during a different time period and potentially subject to uncontrolled environmental variables (e.g., seasonal variation, wildfire smoke, or ambient chemical fluctuations) that were not controlled for. Second, our quantification approach differed in both scope and technique: we used stitched, whole-section raster scanning to comprehensively assess both the dorsal and ventral hippocampus, rather than sampling isolated fields of view. This approach reduces the potential for sampling bias, particularly toward plaque-dense areas, and likely yields more conservative and regionally balanced estimates of plaque burden. It also reflects the true heterogeneity of pathology across the hippocampus, which may mask localized effects of TRAP observed in smaller, targeted analyses.

Nevertheless, plaque counts alone may be an incomplete indicator of functional neuropathology. While total PM particle counts trended higher in the dorsal hippocampus for TRAP-exposed females, these differences did not reach statistical significance, suggesting sex-linked differences in particle accumulation may be subtle or secondary to regional barrier properties. Notably, hyperspectral analysis revealed ventral hippocampal particles exhibited more pronounced red shifts overall, potentially reflecting enhanced biochemical transformation or interaction with the local microenvironment. Within the dorsal hippocampus, particles from TRAP-exposed animals displayed greater spectral red-shifting than those from FA controls, indicating that TRAP exposure alters the chemical signature of retained particles, even in regions with comparable PM counts.

Together, these data suggest TRAP exposure does not uniformly increase brain particle load but rather modulates the biochemical fate of particles in a region- and sex-dependent manner. The dorsal hippocampus represents a site of converging pathology, higher plaque burden, elevated phagocytic microglia, and altered particle chemistry under TRAP conditions, particularly in females, highlighting the intersection of environmental insult and intrinsic neurodegenerative vulnerability.

### Mechanistic Considerations

The above findings suggest several mechanistic hypotheses about how TRAP-derived particles enter and interact with the brain, and why their effects are modulated by region and sex. The regional distribution of particles offers further insight into their route of brain entry. The olfactory system provides a well-established route of access that bypasses the BBB, allowing direct access from the nasal cavity to the olfactory bulb and subsequently to connected regions like the entorhinal cortex and ventral hippocampus.^19^ This pathway has been implicated in early hippocampal and entorhinal cortex pathology in air pollution-exposed animals, including tau accumulation and neuron loss.^20,21^ Yet, we observed the highest particle accumulation in the dorsal hippocampus, particularly in TRAP-exposed females. This pattern is not consistent with a dominant olfactory transport mechanism, which would be expected to preferentially affect more ventral and anterior brain regions. Instead, enrichment of particles in the dorsal hippocampus suggests a hematogenous route of entry,^3,22^ whereby circulating ultrafine particles gain access to the brain parenchyma via compromised BBB integrity.

If BBB disruption facilitates PM translocation from the bloodstream into brain tissue, then regions like dorsal CA1, which may be inherently more vascularized or more sensitive to barrier breakdown, could preferentially accumulate particulate material.^23–25^ Thus, the dorsal-predominant distribution of TRAP particles, alongside sex-specific vulnerability, supports a systemic route of particle delivery to the brain parenchyma rather than retrograde transport via the olfactory nerve.

Once PM_2.5_ gains access to the brain parenchyma, microglia are thought to be the primary immune sentinels that govern their fate.^26,27^ Particle uptake has been extensively studied in microglia both *in vitro* and *in vivo*, with particular focus on PM_2.5_-induced microglial activation and inflammation and subsequent impact on neural function and promotion of neurodegeneration.^27^ However, our data challenge the prevailing assumption that direct microglial sequestration of particles is necessary to induce inflammation. Despite robust microglial phagocytic activity in TRAP-exposed animals, as indicated by elevated CD68 expression in Iba1 immunopositive cells, we observed that PM particles were rarely internalized in microglia or embedded within Aβ plaques. Instead, particles accumulated near, but not within, these cellular and pathological structures, respectively, suggesting a spatially adjacent but non-sequestered interaction. This unexpected pattern implies that while microglia respond to TRAP exposure, they may not effectively capture or internalize particles, especially in chronically inflamed or plaque-rich regions.

One interpretation is that microglia in the TRAP-exposed AD brain may be in a “primed” functionally altered state, but not efficiently phagocytic. Such a state has been previously described for disease-associated microglia (DAMs), which cluster around Aβ but often exhibit impaired debris clearance.^28^

Taken together, our findings suggest that PM-microglia-Aβ plaque interactions are governed more by extracellular crosstalk and biochemical transformation than by direct interactions. This raises the possibility that TRAP-derived particles amplify neuroinflammation and tissue damage by persisting extracellularly in vulnerable regions, interacting with inflammatory mediators, and undergoing surface modifications that influence their optical properties.

While we interpret the red-shifted spectral profiles of particles near plaques and microglia as evidence of *in situ* transformation within pathological tissue microenvironments, an alternate or complementary explanation is that the particles themselves differ in composition at the time of brain entry. TRAP is a heterogeneous mixture, and its component particles can vary by source (e.g., combustion-derived metals vs. tire wear vs. secondary organic aerosols).^29^ Other sources of refractive PM may also give rise to the variable spectral classes of particles measured, such as inhaled nanoplastics.^30^ If different particle classes have varying capacities to penetrate the BBB or persist in neural tissue, then spectral differences may reflect selective deposition or retention of particle types with distinct optical properties. For example, iron-rich or carbonaceous particles exhibit higher refractive indices and specific scattering profiles, which could contribute to the observed red shift independent of biological interaction.

### Limitations

Several limitations of our study should be acknowledged when interpreting these results. First, we utilized a single transgenic rat model of AD (TgF344-AD), which carries familial AD mutations (APP^Swe^ and PS1^ΔE9^) and develops both Aβ plaques and tau tangles with age.^13^ While this model recapitulates many features of human AD, the findings may not generalize to other models or sporadic AD. The exclusive focus on TgF344-AD rats means we cannot discern how normal aging brains (without AD pathology) would respond to the same chronic TRAP exposure, a pertinent comparison given that neuroinflammation in response to pollution can occur even in the absence of underlying amyloidosis.^16,31^

Second, our hyperspectral imaging approach to identify and characterize brain PM has inherent technical limitations. EDF-HSI allows label-free detection of nanoscale particles in tissue, but it cannot definitively determine particle composition or differentiate single particles from small aggregates below the optical resolution limit (∼300-400 nm). We relied on spectral analysis to classify particles, which provides only approximate identification (e.g., classifying a pixel as “particle-like” based on its spectrum). Moreover, the spectral resolution is modest, so different materials can share similar spectra, potentially confounding precise identification. In addition, HSI measures do not directly reveal the chemical nature of particles, nor the exact biomolecules adsorbed onto them.

A third limitation is the scope of brain regions and endpoints analyzed. We concentrated on the hippocampus (dorsal and ventral segments) as a region highly relevant to AD and susceptible to both Aβ plaque and neuroinflammatory changes.^32,33^ However, by doing so, we may have missed effects occurring in other brain areas (particularly the olfactory bulb), other immune pathways (astrocytes), and other cellular features (BBB proteins) impacted by TRAP. Our previous data suggested that the hippocampus had the highest particle deposition, with much lower particle counts in the cortex, thalamus, and cerebellum.^8^ We also primarily assessed end-stage outcomes (after 14 months of exposure); thus, it remains unclear how these changes emerged or whether there were transient spikes or effects that we did not capture earlier in the exposure. Our sample size, while adequate to detect medium-to-large effects, was relatively limited when broken down by sex, region, and other factors (n=6 per sex/exposure subgroup for most measures). This may have constrained our power to detect subtle interactions, for instance, trends toward differences in particle load might have reached significance with a larger cohort. Finally, the study was largely correlative and descriptive in nature. We did not directly test mechanistic pathways; therefore, any inferences made are based on the interpretation of existing literature. Despite these limitations, the convergence of our imaging and spectral data provides a coherent picture that TRAP exposure stimulates a unique pattern of neuroinflammation and particulate deposition in the AD brain.

This study highlights long-term exposure (over a year) to real-world TRAP can alter the particulate landscape in a sex- and region-dependent manner, without necessarily increasing Aβ plaque burden in the brain of a rodent model of AD. The implications for human health are that individuals with prodromal AD or high AD risk (especially females) living in high-pollution areas might experience heightened brain inflammation or accelerated tau pathology even if Aβ plaque accumulation is unaffected. Ultimately, our findings reinforce the concept that environmental factors like air pollution are not just lung and cardiovascular hazards, but also potential contributors to the brain aging and dementia, acting in concert with genetic and sex-specific factors to influence disease trajectories. Taking action to improve air quality could therefore have far-reaching benefits for brain health, particularly in vulnerable populations at risk for AD.

## Materials and Methods

An overview of essential materials and methods is included in this section. Full experimental protocols and materials can be found in the **Supporting Information**, including detailed methods for animal exposure, brain sectioning, immunofluorescent labeling, confocal microscopy, hyperspectral imaging, image processing, particle quantification, and statistical analysis.

### Animals

All experimental protocols involving animals were approved by the UC Davis Institutional Animal Care and Use Committee (IACUC). Male hemizygous TgF344-AD rats were bred with wild-type Fischer 344 females, and offspring were assigned to FA or TRAP exposure groups as described previously.^8,12^ Animals were housed in a vivarium adjacent to a vehicular tunnel to enable real-time exposure to TRAP or filtered air, at concentrations consistent with current air quality standards. After 14 months, animals were euthanized, and brains were harvested for analysis.

### Exposure System and Sample Preparation

The tunnel exposure paradigm and air quality filtration system for the FA control have been described previously.^34,35^ Briefly, air was continuously drawn from an active traffic tunnel and routed unaltered to the TRAP exposure chamber. Control air was filtered using a multi-stage system including activated carbon and high-efficiency particle filters. The hippocampus was dissected post-perfusion and processed into 5 μm cryosections for downstream imaging and analysis.

### Imaging and Quantification

Confocal fluorescence and enhanced darkfield hyperspectral imaging (EDF-HSI) were used to visualize and quantify microglial markers, Aβ plaques, and particulate matter. Antibodies targeting Iba1 and CD68 were used for microglial phenotyping, while Thioflavin S staining identified Aβ plaques. Hyperspectral imaging of unstained tissue enabled detection and spectral profiling of deposited TRAP particles. Fluorescent and hyperspectral images were stitched, registered, and analyzed using a combination of commercial (BioTek Gen5, ENVI) and custom Python-based tools. Statistical analyses were performed in GraphPad Prism using ANOVA with post hoc comparisons as appropriate. Full experimental protocols and details for staining, imaging parameters, spectral processing, ROI selection, image analysis, and software workflows can be found in the **Supporting Information.**

## Author Contributions

The manuscript was written through contributions of all authors. All authors have approved the final version of the manuscript. **Hannah J. O’Toole:** Conceptualization, Methodology; Investigation, Formal analysis, Visualization, Writing – original draft; Writing – review & editing. **Anchaleena James:** Methodology, Investigation, Data curation, Visualization; **Nathifa Nasim:** Methodology, Investigation, Writing – review & editing; **Dustin J. Hadley:** Methodology, Data curation, Formal analysis. **Elizabeth J. Hale:** Investigation, Writing – review & editing. **Qing He:** Methodology, Writing – review & editing. **Keith J. Bein:** Conceptualization, Methodology, Resources. **Anthony Venezuela:** Methodology, Resources, Data collection. **Tatu Rojalin:** Conceptualization, Methodology. **Brittany N. Dugger:** Conceptualization; Writing – review & editing. **Anthony S. Wexler:** Conceptualization, Supervision, Writing – review & editing. **Pamela J. Lein:** Conceptualization, Supervision, Funding acquisition, Project administration, Writing – review & editing; **Randy P. Carney:** Conceptualization, Supervision, Funding acquisition, Project administration, Methodology, Investigation, Formal analysis, Writing – original draft; Writing – review & editing.

## Acknowledgements

This work was financially supported by the National Institutes of Health (NIH) by Award Numbers R01 NS13065 and R01 CA241666, and the National Science Foundation (NSF) Project Number 2238995. H.J.O. acknowledges T32 GM136597. The funders had no roles in the study design, data collection, data analysis, decision to publish, or preparation of the manuscript. The content of this work is solely that of the authors and does not necessarily represent the views of the funding parties. Components of figures in this work were created using or inspired by BioRender.com. EDF-HSI imaging was conducted using the CAMI core facility at the UC Davis Center for Health and the Environment. The CytoViva was funded by NIH S10 OD028584. Thanks to CAMI core staff member Morgan Domanico, Ph.D., for her assistance with instrumentation and training.

## Conflict of Interest Statement

The authors declare no conflicts of interest.

## Notes

### Competing Interest Statement

The authors have declared no competing interest.

## References

(1) Costa, L. G. Traffic-Related Air Pollution and Neurodegenerative Diseases: Epidemiological and Experimental Evidence, and Potential Underlying Mechanisms. In Advances in Neurotoxicology; Elsevier, 2017; Vol. 1, pp 1–46. 10.1016/bs.ant.2017.07.001.

(2) Blanco, M. N.; Shaffer, R. M.; Li, G.; Adar, S. D.; Carone, M.; Szpiro, A. A.; Kaufman, J. D.; Larson, T. V.; Hajat, A.; Larson, E. B.; Crane, P. K.; Sheppard, L. Traffic-Related Air Pollution and Dementia Incidence in the Adult Changes in Thought Study. Environment International 2024, 183, 108418. 10.1016/j.envint.2024.108418.

(3) Park, H.; Armstrong, M.; Gorin, F.; Lein, P. Air Pollution as an Environmental Risk Factor for Alzheimer’s Disease and Related Dementias. MRAJ 2024, 12 (10). 10.18103/mra.v12i10.5825.

(4) Weuve, J.; Bennett, E. E.; Ranker, L.; Gianattasio, K. Z.; Pedde, M.; Adar, S. D.; Yanosky, J. D.; Power, M. C. Exposure to Air Pollution in Relation to Risk of Dementia and Related Outcomes: An Updated Systematic Review of the Epidemiological Literature. Environ Health Perspect 2021, 129 (9), 096001. 10.1289/EHP8716.

(5) Win-Shwe, T.-T.; Fujimaki, H. Nanoparticles and Neurotoxicity. IJMS 2011, 12 (9), 6267–6280. 10.3390/ijms12096267.

(6) Lundborg, M.; Johard, U.; Låstbom, L.; Gerde, P.; Camner, P. Human Alveolar Macrophage Phagocytic Function Is Impaired by Aggregates of Ultrafine Carbon Particles. Environmental Research 2001, 86 (3), 244–253. 10.1006/enrs.2001.4269.

(7) Li, N.; Sioutas, C.; Cho, A.; Schmitz, D.; Misra, C.; Sempf, J.; Wang, M.; Oberley, T.; Froines, J.; Nel, A. Ultrafine Particulate Pollutants Induce Oxidative Stress and Mitochondrial Damage. Environ Health Perspect 2003, 111 (4), 455–460. 10.1289/ehp.6000.

(8) Patten, K. T.; Valenzuela, A. E.; Wallis, C.; Berg, E. L.; Silverman, J. L.; Bein, K. J.; Wexler, A. S.; Lein, P. J. The Effects of Chronic Exposure to Ambient Traffic-Related Air Pollution on Alzheimer’s Disease Phenotypes in Wildtype and Genetically Predisposed Male and Female Rats. Environ Health Perspect 2021, 129 (5), 057005. 10.1289/EHP8905.

(9) Jayaraj, R. L.; Rodriguez, E. A.; Wang, Y.; Block, M. L. Outdoor Ambient Air Pollution and Neurodegenerative Diseases: The Neuroinflammation Hypothesis. Curr Envir Health Rpt 2017, 4 (2), 166–179. 10.1007/s40572-017-0142-3.

(10) Bein, K. J.; Wallis, C. D.; Silverman, J. L.; Lein, P. J.; Wexler, A. S. Emulating Near-Roadway Exposure to Traffic-Related Air Pollution via Real-Time Emissions from a Major Freeway Tunnel System. Environ. Sci. Technol. 2022, 56 (11), 7083–7095. 10.1021/acs.est.1c07047.

11. California Air Resources Board. Statewide SIPs: 9 Μg/M^3^ PM. ; Webpage; California Air Resources Board, 2024. https://ww2.arb.ca.gov/our-work/programs/california-state-implementation-plans/statewide-efforts/sips-9-mgm3-pm2-5 (accessed 2025-06-25).

(12) Cohen, R. M.; Rezai-Zadeh, K.; Weitz, T. M.; Rentsendorj, A.; Gate, D.; Spivak, I.; Bholat, Y.; Vasilevko, V.; Glabe, C. G.; Breunig, J. J.; Rakic, P.; Davtyan, H.; Agadjanyan, M. G.; Kepe, V.; Barrio, J. R.; Bannykh, S.; Szekely, C. A.; Pechnick, R. N.; Town, T. A Transgenic Alzheimer Rat with Plaques, Tau Pathology, Behavioral Impairment, Oligomeric Aβ, and Frank Neuronal Loss. J. Neurosci. 2013, 33 (15), 6245–6256. 10.1523/JNEUROSCI.3672-12.2013.

(13) Nataraj, A.; Blahna, K.; Ježek, K. Insights From TgF344-AD, a Double Transgenic Rat Model in Alzheimer’s Disease Research. Physiol Res 2025, No. 1/2025, 1–17. 10.33549/physiolres.935464.

(14) McInnes, L.; Healy, J.; Saul, N.; Großberger, L. UMAP: Uniform Manifold Approximation and Projection. JOSS 2018, 3 (29), 861. 10.21105/joss.00861.

(15) O’Piela, D. R.; Durisek, G. R.; Escobar, Y.-N. H.; Mackos, A. R.; Wold, L. E. Particulate Matter and Alzheimer’s Disease: An Intimate Connection. Trends in Molecular Medicine 2022, 28 (9), 770–780. 10.1016/j.molmed.2022.06.004.

(16) Patten, K. T.; Valenzuela, A. E.; Wallis, C.; Harvey, D. J.; Bein, K. J.; Wexler, A. S.; Gorin, F. A.; Lein, P. J. Hippocampal but Not Serum Cytokine Levels Are Altered by Traffic-Related Air Pollution in TgF344-AD and Wildtype Fischer 344 Rats in a Sex- and Age-Dependent Manner. Front. Cell. Neurosci. 2022, 16, 861733. 10.3389/fncel.2022.861733.

(17) Cacciottolo, M.; Wang, X.; Driscoll, I.; Woodward, N.; Saffari, A.; Reyes, J.; Serre, M. L.; Vizuete, W.; Sioutas, C.; Morgan, T. E.; Gatz, M.; Chui, H. C.; Shumaker, S. A.; Resnick, S. M.; Espeland, M. A.; Finch, C. E.; Chen, J. C. Particulate Air Pollutants, APOE Alleles and Their Contributions to Cognitive Impairment in Older Women and to Amyloidogenesis in Experimental Models. Transl Psychiatry 2017, 7 (1), e1022–e1022. 10.1038/tp.2016.280.

(18) Mahmoudi, M.; Landry, M. P.; Moore, A.; Coreas, R. The Protein Corona from Nanomedicine to Environmental Science. Nat Rev Mater 2023, 8 (7), 422–438. 10.1038/s41578-023-00552-2.

(19) Ji, X.; Liu, R.; Guo, J.; Li, Y.; Cheng, W.; Pang, Y.; Zheng, Y.; Zhang, R.; Tang, J. Olfactory Bulb Microglia Activation Mediated Neuronal Death in Real-Ambient Particulate Matter Exposure Mice with Depression-like Behaviors. Science of The Total Environment 2022, 821, 153456. 10.1016/j.scitotenv.2022.153456.

(20) Chuang, H.-C.; Chen, H.-C.; Chai, P.-J.; Liao, H.-T.; Wu, C.-F.; Chen, C.-L.; Jhan, M.-K.; Hsieh, H.-I.; Wu, K.-Y.; Chen, T.-F.; Cheng, T.-J. Neuropathology Changed by 3- and 6-Months Low-Level PM2.5 Inhalation Exposure in Spontaneously Hypertensive Rats. Part Fibre Toxicol 2020, 17 (1), 59. 10.1186/s12989-020-00388-6.

(21) Lee, S.-H.; Chen, Y.-H.; Chien, C.-C.; Yan, Y.-H.; Chen, H.-C.; Chuang, H.-C.; Hsieh, H.-I.; Cho, K.-H.; Kuo, L.-W.; Chou, C. C.-K.; Chiu, M.-J.; Tee, B. L.; Chen, T.-F.; Cheng, T.-J. Three Month Inhalation Exposure to Low-Level PM2.5 Induced Brain Toxicity in an Alzheimer’s Disease Mouse Model. PLoS ONE 2021, 16 (8), e0254587. 10.1371/journal.pone.0254587.

(22) Miller, M. R.; Raftis, J. B.; Langrish, J. P.; McLean, S. G.; Samutrtai, P.; Connell, S. P.; Wilson, S.; Vesey, A. T.; Fokkens, P. H. B.; Boere, A. J. F.; Krystek, P.; Campbell, C. J.; Hadoke, P. W. F.; Donaldson, K.; Cassee, F. R.; Newby, D. E.; Duffin, R.; Mills, N. L. Inhaled Nanoparticles Accumulate at Sites of Vascular Disease. ACS Nano 2017, 11 (5), 4542–4552. 10.1021/acsnano.6b08551.

(23) Montagne, A.; Barnes, S. R.; Sweeney, M. D.; Halliday, M. R.; Sagare, A. P.; Zhao, Z.; Toga, A. W.; Jacobs, R. E.; Liu, C. Y.; Amezcua, L.; Harrington, M. G.; Chui, H. C.; Law, M.; Zlokovic, B. V. Blood-Brain Barrier Breakdown in the Aging Human Hippocampus. Neuron 2015, 85 (2), 296–302. 10.1016/j.neuron.2014.12.032.

(24) Joo, I. L.; Lai, A. Y.; Bazzigaluppi, P.; Koletar, M. M.; Dorr, A.; Brown, M. E.; Thomason, L. A. M.; Sled, J. G.; McLaurin, J.; Stefanovic, B. Early Neurovascular Dysfunction in a Transgenic Rat Model of Alzheimer’s Disease. Sci Rep 2017, 7 (1), 46427. 10.1038/srep46427.

(25) Smith, N. M.; Gachulincova, I.; Ho, D.; Bailey, C.; Bartlett, C. A.; Norret, M.; Murphy, J.; Buckley, A.; Rigby, P. J.; House, M. J.; St. Pierre, T.; Fitzgerald, M.; Iyer, K. S.; Dunlop, S. A. An Unexpected Transient Breakdown of the Blood Brain Barrier Triggers Passage of Large Intravenously Administered Nanoparticles. Sci Rep 2016, 6 (1), 22595. 10.1038/srep22595.

(26) Zhang, L.; Xu, F.; Yang, Y.; Yang, L.; Wu, Q.; Sun, H.; An, Z.; Li, J.; Wu, H.; Song, J.; Wu, W. PM2.5 Exposure Upregulates pro-Inflammatory Protein Expression in Human Microglial Cells via Oxidant Stress and TLR4/NF-κB Pathway. Ecotoxicology and Environmental Safety 2024, 277, 116386. 10.1016/j.ecoenv.2024.116386.

(27) Ishihara, Y.; Tanaka, M.; Nezu, N.; Ishihara, N.; Oguro, A.; Vogel, C. F. A. Pathways to the Brain: Impact of Fine Particulate Matter Components on the Central Nervous System. Antioxidants 2025, 14 (6), 730. 10.3390/antiox14060730.

(28) Sun, Z.; Zhang, X.; So, K.-F.; Jiang, W.; Chiu, K. Targeting Microglia in Alzheimer’s Disease: Pathogenesis and Potential Therapeutic Strategies. Biomolecules 2024, 14 (7), 833. 10.3390/biom14070833.

(29) Boogaard, H.; Patton, A. P.; Atkinson, R. W.; Brook, J. R.; Chang, H. H.; Crouse, D. L.; Fussell, J. C.; Hoek, G.; Hoffmann, B.; Kappeler, R.; Kutlar Joss, M.; Ondras, M.; Sagiv, S. K.; Samoli, E.; Shaikh, R.; Smargiassi, A.; Szpiro, A. A.; Van Vliet, E. D. S.; Vienneau, D.; Weuve, J.; Lurmann, F. W.; Forastiere, F. Long-Term Exposure to Traffic-Related Air Pollution and Selected Health Outcomes: A Systematic Review and Meta-Analysis. Environment International 2022, 164, 107262. 10.1016/j.envint.2022.107262.

(30) Liu, X.; Zhao, Y.; Dou, J.; Hou, Q.; Cheng, J.; Jiang, X. Bioeffects of Inhaled Nanoplastics on Neurons and Alteration of Animal Behaviors through Deposition in the Brain. Nano Lett. 2022, 22 (3), 1091–1099. 10.1021/acs.nanolett.1c04184.

(31) Zhang, X.; Yin, X.; Zhang, J.; Li, A.; Gong, H.; Luo, Q.; Zhang, H.; Gao, Z.; Jiang, H. High-Resolution Mapping of Brain Vasculature and Its Impairment in the Hippocampus of Alzheimer’s Disease Mice. National Science Review 2019, 6 (6), 1223–1238. 10.1093/nsr/nwz124.

(32) Furcila, D.; Domínguez-Álvaro, M.; DeFelipe, J.; Alonso-Nanclares, L. Subregional Density of Neurons, Neurofibrillary Tangles and Amyloid Plaques in the Hippocampus of Patients With Alzheimer’s Disease. Front. Neuroanat. 2019, 13, 99. 10.3389/fnana.2019.00099.

(33) Marlatt, M. W.; Bauer, J.; Aronica, E.; Van Haastert, E. S.; Hoozemans, J. J. M.; Joels, M.; Lucassen, P. J. Proliferation in the Alzheimer Hippocampus Is Due to Microglia, Not Astroglia, and Occurs at Sites of Amyloid Deposition. Neural Plasticity 2014, 2014, 1–12. 10.1155/2014/693851.

(34) Edwards, S.; Zhao, G.; Tran, J.; Patten, K. T.; Valenzuela, A.; Wallis, C.; Bein, K. J.; Wexler, A. S.; Lein, P. J.; Rao, X. Pathological Cardiopulmonary Evaluation of Rats Chronically Exposed to Traffic-Related Air Pollution. Environ Health Perspect 2020, 128 (12), 127003. 10.1289/EHP7045.

(35) Patten, K. T.; González, E. A.; Valenzuela, A.; Berg, E.; Wallis, C.; Garbow, J. R.; Silverman, J. L.; Bein, K. J.; Wexler, A. S.; Lein, P. J. Effects of Early Life Exposure to Traffic-Related Air Pollution on Brain Development in Juvenile Sprague-Dawley Rats. Transl Psychiatry 2020, 10 (1), 166. 10.1038/s41398-020-0845-3.

