## Supporting Information for "Spatial and spectral mapping of traffic-related air pollution (TRAP) nanoparticles in relation to plaques and inflammatory markers in an Alzheimer disease model"

**Supporting Methods**
**Animals:** Animal procedures were performed in compliance with protocols approved by the University of California Davis (UC Davis) Institutional Animal Care and Use Committee (IACUC), with the goal of minimizing pain and suffering. A breeding colony of TGF344-AD rats was established using male hemizygous TgF344-AD rats acquired from Emory University and female WT Fischer (F344) rats purchased from Charles River Laboratories (Charles River Laboratories International Inc.; Wilmington, MA, USA), as previously described.^1^ Pups were randomly assigned to different exposure groups (FA or TRAP) using a random number generator and were tattooed with coded identification numbers using nontoxic animal tattoo ink (Ketchum Manufacturing Inc.; Brockville, ON, Canada) at postnatal (P) day 2 (P2). Pups were genotyped at P8 as previously described^2^ and weaned at P21. P28 animals were transported to the tunnel exposure facility vivarium approved by the UC Davis IACUC, as previously described^3^. The vivarium was maintained under controlled environmental conditions (20-26°C, 12:12 light:dark cycle) with food (Envigo Global 18% Protein Rodent Diet; Inotiv, Inc.; West Lafayette, IN, USA) and filtered water provided ad libitum. After 14 months of exposure, animals were transported back to the UC Davis campus, housed in an IACUC-approved rodent vivarium at the UC Davis Center for Health and the Environment (CHE), and euthanized 18 h later to collect tissues. The same husbandry protocols and conditions of the tunnel facility were maintained at CHE, except for ambient traffic noise and vibration, and exposure to standard vivarium air.

**Exposure and TRAP Characterization:** The tunnel exposure facility housed animals within exposure chambers drawing on TRAP or FA. TRAP animals were continuously exposed to air delivered unchanged in real-time from a tunnel bore with both light- and heavy-duty vehicular traffic. TRAP was characterized as previously described.^3,4^ Multiple emission control technologies were used for the FA control exposure group. FA control animals were exposed to ambient air collected from a storage shed immediately adjacent to the vivarium which was filtered by coarse filtration to remove debris and dust, inline activated carbon for volatile organic compounds, barium oxide-based catalytic converters for removing nitrogen oxides, and a 6-port ultrahigh-efficiency particle filtering system for removing ultrafine, fine, and coarse PM^3,4^ prior to exposure to the FA control animals. The airflow rate of the exposure chambers was 35 ft^3^/min, as per IACUC specifications.

**Sample Preparation:** Upon perfusion, the right hemisphere of the brain was cut into 2-mm coronal blocks and fixed in 4% wt/vol paraformaldehyde (PFA; Sigma Chemical) in 0.1M phosphate buffer (pH 7.2, 10 mM sodium phosphate dibasic, 1.8 mM potassium phosphate monobasic) for 24 h at 4°C. After 3×5 min washes in PBS, blocks were sunk in 30% wt/vol sucrose (ThermoFisher Scientific) in PBS for 1 week. Blocks were then embedded in Optimal Cutting Temperature (OCT) compound (ThermoFisher Scientific) and flash frozen in a bath of dry ice and methanol. Frozen blocks were then cryosectioned into 5 μm-thick sections. Targeted regions included the dorsal hippocampus (bregma -3.3 to -4.2) and one ventral hippocampal section (bregma -4.7 to -5.6), confirmed using *The Rat Brain in Stereotaxic Coordinates*.

**Immunofluorescent Labeling:** Fluorescent immunohistochemistry (IHC) was performed on 5µm thick sections placed on Fisherbrand™ Superfrost™ Plus Regular Microscope Slides (Cat. No. 12-550-15; Thermo Fisher Scientific; Waltham, MA, USA). For each stain, n=6 rats per group per sex were analyzed, each with two slides with two sections per slide (technical replicates). Antigen retrieval was done by heating slides in sodium citrate buffer (10 mM, pH 6.0) at 90°C for 30 min. Slides were then washed 3x in 0.03% PBST for 5 min. Slides were then incubated with primary antibodies against microglial markers. Briefly, they were incubated with mouse anti-rat CD68 at 1:200 (Cat. No. MCA341R; Bio-Rad Laboratories; Hercules, CA, USA) and rabbit anti-rat Iba1 at 1:1000 (Cat. No. 019-19741; FUJIFILM Wako Chemicals USA, Corp.; Richmond, VA, USA) in 10% BSA and 0.03% PBST overnight at 4°C. Slides were then washed 3x in 1X PBS for 5 min, then incubated with fluorescent secondary antibodies. Briefly AlexaFluor 488 Goat anti-mouse IgG1 at 1:1000 (Cat. No. A21121; Invitrogen; Thermo Fisher Scientific; Waltham, MA, USA) and AlexaFluor 568 Goat anti-rabbit IgG at 1:1000 (Cat. No. A11036; Invitrogen; Thermo Fisher Scientific; Waltham, MA, USA) were incubated in 0.03% PBST for 1 hour at room temperature. For negative controls, a blocking buffer was used in place of the primary antibody. Slides were washed with PBS and mounted in ProLong™ Gold Antifade Mountant with DNA stain 4′,6-diamidino-2-phenylindole (DAPI) (Cat. No. P3691; Invitrogen; Thermo Fisher Scientific; Waltham, MA, USA) and Fisherbrand™ Superslip™ Coverslips (Cat. No. 12-541-057; Thermo Fisher Scientific; Waltham, MA, USA) were placed on top of the slides.

**Thioflavin S Staining:** The second slide with subsequent sections were stained for Thioflavin S (ThioS) to visualize Aβ-plaques; fixed brain sections were stained with 0.02% wt/vol Thioflavin S (Sigma Chemical) in water for 8 min and then de-stained with 50% vol/vol ethanol (Thermo Fisher Scientific; Waltham, MA, USA) for 3×1 min. Slides were mounted in ProLong™ Gold without DAPI (Thermo Fisher Scientific; Waltham, MA, USA) and coverslipped with Fisherbrand™ Superslip™ Coverslips (Cat. No. 12-541-057; Thermo Fisher Scientific; Waltham, MA, USA).

**Confocal Fluorescence Microscopy:** Fluorescent images were acquired via confocal microscopy using the Agilent BioTek Cytation C10 confocal imaging reader (Agilent Technologies, Inc.; Santa Clara, CA, USA). Both the ThioS-stained slides and Iba1/CD68/DAPI slides (preparations described above) were first imaged using the Cytation C10 widefield brightfield at 4x to localize the hippocampal region of interest (either ventral or dorsal, (n=24 total regions: n=12 dorsal, n=12 ventral; n=3 of each sex and exposure condition)) in each mounted coronal rat brain section on both the ThioS and Iba1/CD68/DAPI slides. Once visualized at 4x, a spatial montage was selected to generate an ROI to take confocal fluorescence images at 20x PL FL Phase using the 6 μm disk. For ThioS-stained slides, images were obtained using the DAPI channel (DAPI Confocal Imaging Cube, Part No. 1945103) with 150 ms integration time and 28 camera gain. The DAPI channel was used to set the Z-stack positions for these slides. For the Iba1/CD68/DAPI slides, images were obtained using the TRITC channel (TRITC Confocal Imaging Cube, Part No. 1945106; ex: 556/20 nm, em: 600/37 nm, cut-off: 573 nm; 150 ms integration time, 31.1 camera gain), GFP channel (GFP Confocal Imaging Cube, Part No. 1945104; ex: 472/30 nm, em: 520/35 nm, cut-off: 495 nm; 150 ms integration time, 26 camera gain), and DAPI channel to visualize cell nuclei (DAPI Confocal Imaging Cube, Part No. 1945103; ex: 390/40 nm, em: 442/42 nm, cut-off: 414 nm; 400 ms integration time, 31.5 camera gain), respectively. Depending on the floor and top Z-positions of each section, a different number of Z-slices were acquired for each slide. Initial image processing was performed within the BioTek Gen5 Image Prime software (Version 3.16; Agilent Technologies, Inc.; Santa Clara, CA, USA) that accompanies the Cytation C10 confocal imaging reader. Briefly, images were deconvolved, underwent Z-projection based on focus stacking, and were stitched (linear blend fusion method, filling gaps between montage tiles with local background) based on the TRITC channel for the Iba1/CD68/DAPI slides, and the DAPI channel for the ThioS slides. To keep image export <1 GB, some images were downsized to the suggested downsize percentage within the BioTek Gen5 Image Prime software.

**Enhanced-Darkfield Hyperspectral Imaging:** Enhanced-dark field hyperspectral imaging (EDF-HSI) was performed using the CytoViva Enhanced Darkfield Illuminator Microscope configuration (CytoViva, Inc.; Auburn, AL, USA) at the UC Davis CAMI core facility to visualize particulate matter (PM) and Aβ-plaque formations without the need for labeling. By integrating EDF imaging with confocal microscopy, we were able to accurately assess the location and density of PM in relation to the surrounding tissue and neuroanatomy. Confocal fluorescence images were used as guides to navigate the slides for EDF-HSI imaging. The same slides as used for confocal fluorescence were used for EDF-HSI. The CytoViva EDF is equipped with a motorized stage, enabling mapping and precise navigation around the slides. A 10x objective (Olympus PLN Plan Achromat 10x Microscope Objective, N.A. 0.25, W.D. 10.6, Part. No. 1-U2B233; Olympus Corporation; Tokyo, Japan) was utilized to identify the hippocampal region of interest and take note of any landmarks within the tissue. Landmarks were manually digitally annotated to enable navigation during imaging. For each imaged hippocampal region on each slide, an origin was selected at a tissue landmark, the automatic stage was zeroed, and coordinates were monitored to determine whereabouts during manual imaging. Enhanced-darkfield images were acquired at 40x using the Olympus LUCPlanFL N 40x objective lens with correction collar (N.A. 0.60, Olympus Corporation; Tokyo, Japan; using Ocular USB software (Ocular Advanced Scientific Camera Control Version 2.0.1.496 (64-bit), PVCAM Version 3.6.7; Teledyne Vision Solutions; Tucson, AZ, USA). Following EDF image acquisition, hyperspectral images were acquired at the exact same spots as EDF images utilizing Micro-Manager software (Micro-Manager MMStudio Version 1.4.22).^5^ HSI images were acquired using the Photometrics Evolve 512 series EMCCD camera (Teledyne Vision Solutions; Tucson, AZ, USA) at 512 x 512 pixels. All HSI images were acquired at 40x with a 500 ms integration time, high Pre-Gain, “flip horizontal” checked (to match the orientation of the eyepiece) and an EM gain of 5.
**Enhanced-darkfield Widefield Stitched Images:** For the two dorsal hippocampal regions in **Fig. 2A,D**, individual EDF images acquired at 40x were stitched to create a widefield visualization in darkfield. Each individual field of view (FOV) measured approximately 200 µm × 200 µm. To generate whole-section hyperspectral maps, individual tile images were stitched using a custom Python script (Python v3.13.5, utilizing the open-source libraries tifffile, numpy, and opencv) that calculated tile offsets based on microscope acquisition coordinates or encoded filenames. Each tile was assigned a grid position based on a “column, row” naming convention. Overlapping regions were blended using a cosine ramp feathering algorithm to smooth transitions and reduce edge artifacts. Spatial drift across rows and columns was corrected by applying fixed pixel offsets. After all tiles were placed on a shared canvas, pixel intensities were normalized using accumulated blending weights. The final stitched image was cropped to remove empty canvas borders and saved as a single TIFF file.

**Analysis of particle distribution:** To quantify particle burden across hippocampal sections, stitched darkfield hyperspectral images were subdivided into individual tiles based on acquisition coordinates. For each tile, bright spots corresponding to putative UFPM were detected using intensity thresholding and binary contour detection, using a custom Python script (v3.13.5, utilizing the open-source library opencv). Particle counts were overlaid on each image and visually confirmed using a custom interactive GUI that allowed manual adjustment of automated counts. This hybrid approach ensured robust detection while enabling user correction for false positives or overlapping particles. Counts were aggregated across tiles and spatially mapped to reconstruct the distribution of particles across the full brain section. Final data were exported as CSV files for statistical comparison across experimental groups.

**EDF-HSI images for particle analysis:** Particles were identified using EDF microscopy at 10x to visualize landmarks in the surrounding tissue. A total of 24 animals were analyzed (n=6 per group: Female-FA, Female-TRAP, Male-FA, Male-TRAP). Two different areas of the chosen hippocampal tissue regions (dorsal or ventral) were imaged systematically for particle counting. One area was in a region without plaques (ThioS^-^), selected by comparison with the ThioS confocal images, and the other area was a region with Iba1+ markers, selected via the GFP channel confocal images. Plaques were always present within Iba1+ regions, as previously observed.^6^ After establishing an origin at a tissue landmark to match the confocal images of the tissue section, the stage was zeroed and seven Iba1+ regions were randomly selected per hippocampal region per sex per exposure condition and imaged at 40x with EDF using the Ocular USB software. Coordinates for each imaging FOV spot were recorded and annotated on the corresponding GFP channel confocal image. HSI images were acquired at the same coordinates (parameters explained above). For plaque-negative regions, an area of the ThioS-stained hippocampal region of interest with no ThioS signal was selected, and eight FOVs were imaged at 40x in 200 µm FOV intervals with the same procedure as above. Corresponding HSI images at these spots were then collected using identical parameters.

Most animals were assessed in a single hippocampal region (dorsal or ventral), dependent on tissue availability. A small number of animals were analyzed in both regions. For these animals, dorsal and ventral hippocampi were treated as independent data points, as sections were processed, imaged, and quantified separately. For each region, the particle counts from the 15 fields of view (Iba^+^ and ThioS^-^ ) were averaged to yield a single particle count per hippocampal region per animal (**Fig. 2G**). One animal from the Female-TRAP group was excluded in particle counting due to tissue thickness interfering with EDF/HSI acquisition, resulting in n=2 for that condition in the final dataset (**Fig. 2G**). Quantification and statistical analysis were performed at the level of individual hippocampal regions.

**HSI Image Analysis:** HSI analysis was performed using ENVI 4.8 (NV5 Geospatial Software; Broomfield, CO, USA) with CytoViva’s EMCCD plugin. HSI images were processed in ENVI 4.8 to smooth spectra, filter and select particle ROIs, and extract spectral information at selected ROI pixels. HSI images were first smoothed using the Savitsky-Golay Curve Fit Smoothing function within the CytoViva Analysis tab. In this method, noise is removed from the spectrum by fitting curves over segments of the spectrum. As suggested by CytoViva’s manual, default settings (Width: 33, Degree:2) for this feature were used to best retain the original curve amplitude. Particles were then initially filtered using the CytoViva Particle Analysis Particle Filter tool. The Background Filtering Method “Max” was selected, with parameters for "Max must exceed" (intensity counts) and "Valid Data Max" adjusted for each image to adequately select the particles within the image from the background tissue. If an image FOV included a plaque, an ROI was selected on the plaque region to see the maximum spectral intensity at that region, and this value was used as the “Max must exceed” parameter. Plaques appear as brighter regions of the background tissue. No filtering was done for size. Following Particle Filtering, selected particles were reviewed in the Particle Filter Review Toolbox to remove erroneous particle ROI selections. These selected particles were exported to ROI files, then any manual particle selection for any particles missed by the CytoViva Particle Filter was performed within the ROI tool. Once all particles within the FOV were selected, the ROI file was converted to a spectral library file (SLF), containing both mean spectra (representing the average spectra of all pixels within each ROI or particle) and all spectra (containing individual pixel data). Each SLF was then imported into the Spectral Angle Mapper (SAM) tool, where mean spectra for each particle were plotted to analyze their spectral properties. The mean spectra SLF was exported to an ASCII file for downstream analysis in Python. Particle counts (selected ROIs) for each FOV for each imaged hippocampal region were logged in Microsoft Excel (Microsoft; Redmond, WA, USA). These counts are also available within the metadata.

**HSI spectral analysis:** Spectral data were extracted from particle-associated regions across hippocampal sections using ASCII export from the hyperspectral imaging system. Each particle’s full reflectance spectrum was loaded into a custom Python (v3.13.5) pipeline, which parsed sample metadata (sex, exposure group, region, proximity to plaque) and aggregated spectra across multiple animals. Data were standardized and subjected to dimensionality reduction using principal component analysis (PCA) and uniform manifold approximation and projection (UMAP) to visualize spectral diversity and group clustering. Spectral comparisons were performed between predefined groups (e.g., near- vs. far-from-plaque) using unpaired t-tests at each wavelength, followed by Benjamini–Hochberg false discovery rate correction. Wavelengths with statistically significant differences were reported, and spectral features most correlated with dimensionality-reduced coordinates were identified using Pearson correlation and random forest regression. Group-specific average spectra were visualized with standard deviation bands to assess variability. All analyses were performed using open-source libraries including numpy, scikit-learn, umap-learn, statsmodels, and seaborn.

**ROI selection for phagocytic microglia and Aβ-plaques:** CD68 and ThioS stained confocal images were analyzed using Image J FIJI^7^ to quantify phagocytic microglia and Aβ plaques. Both dorsal and ventral hippocampal regions from both sexes across both exposure conditions were analyzed. ROIs were created using the freehand selection tool to isolate the hippocampus region and were thresholded (CD68: 0.05% for all; ThioS: 0.05% for ventral and 0.10% for dorsal regions) using the intermodes method with a dark background to better visualize the partition of the image.

Aβ-plaques and CD68+ cells were analyzed via the Analyze Particles tool with size (CD68: 10–∞ pixels; ThioS: 42–∞ pixels) and circularity (0–1.0) parameters. Bare outlines were displayed, and results were summarized, excluding regions on edges. The resultant data sets were saved and used for statistical analysis. Original images were overlaid with the output ROIs and saved as flattened RGB images. For CD68 analysis, the overlay was merged with Iba1 (red) and DAPI (blue) channels using the Merge Channels tool. Composite images were flattened and saved for further analysis.

**Statistical Analyses:** Statistical analyses were performed using GraphPad Prism 10 software (version 10.4.1; GraphPad, San Diego, CA, USA). Data is presented as scatter dot plots with a line at the mean ± standard deviation unless otherwise noted. Family-wise alpha threshold and confidence level was 0.05 (95% confidence interval) for all multiple comparisons tests, using GraphPad’s GP p value style. All significant differences (p<0.05) are reported pair-wise, denoted with an asterisk (*). For all analyses, we first performed three-way analysis of variance (ANOVA) to assess whether Aβ-plaque counts, CD68+ counts, or particle counts (respectively) were influenced by TRAP exposure, sex, or hippocampal region (dorsal versus ventral). If we found a specific effect of the interaction of these factors, either post hoc analysis or data consolidation into two-way ANOVA was performed, followed by post hoc analysis (Tukey’s multiple comparisons test). Aβ-plaque counts (**Fig. 3**) were first normalized to pixel area within Microsoft Excel (Microsoft; Redmond, WA, USA), to account for the varied area of dorsal versus ventral hippocampal regions. Ordinary two-way ANOVA with post hoc Tukey’s multiple comparisons tests was performed for each analysis. Similar to above, CD68+ counts (**Fig. 4**) were first normalized to pixel area within Microsoft Excel (Microsoft; Redmond, WA, USA), so as to account for the varied area of dorsal versus ventral hippocampal regions. To assess particle counts as related to regions of AD pathology (**Fig. 5A**), particle counts (consolidated by combining counts from hippocampal region (dorsal v. ventral) and sex) from the two exposure groups (FA v. TRAP) were assessed based on their number in no-plaque (ThioS^-^) regions versus Iba1^+^ regions using ordinary two-way ANOVA with post hoc Tukey’s multiple comparisons test. However, no significant difference in interaction or main effects were identified.

**Supporting Figures**

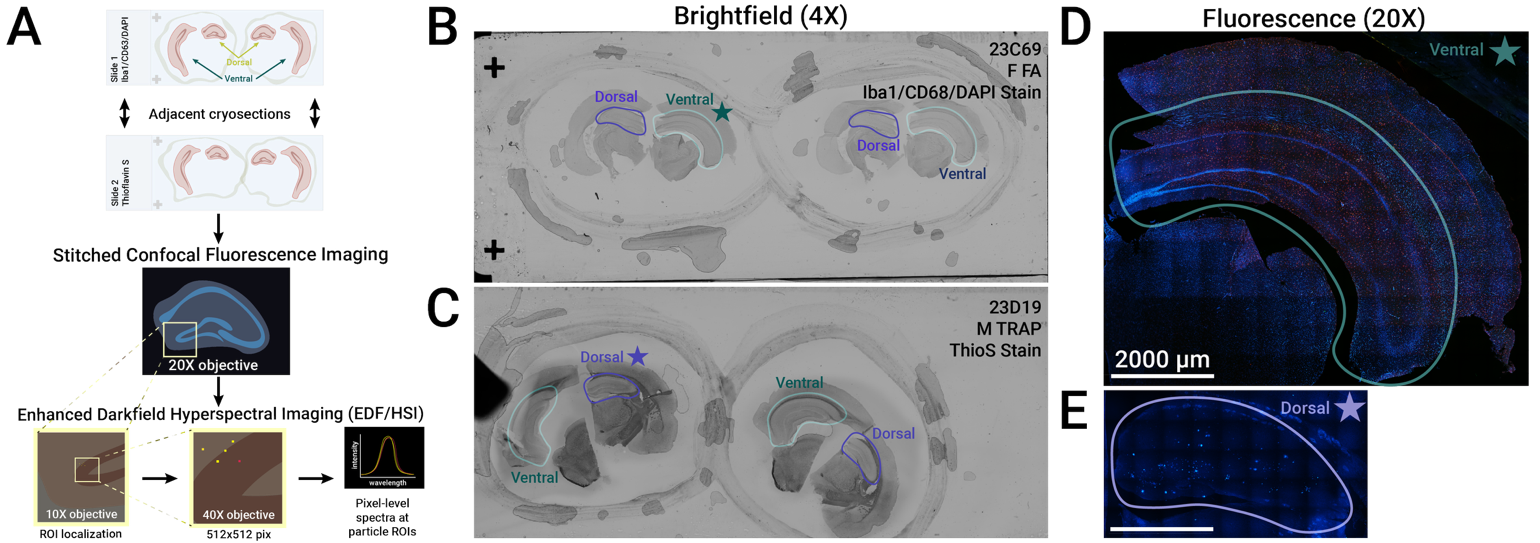

**Supplementary Figure S1. Annotation and determination of dorsal and ventral hippocampal regions for confocal fluorescence and enhanced darkfield hyperspectral imaging analysis.** Two slides per rat were prepared, each containing hippocampal sections including the dorsal and ventral hippocampus. Each slide contained two adjacent cryosections, totaling four adjacent cryosections per rat. One slide was immunostained against Iba1 and CD68 and mounted with DAPI-containing mounting medium (Iba1/CD68/DAPI), and the other slide was stained for Thioflavin S (ThioS) (A). Both slides were imaged via confocal fluorescence microscopy utilizing the Cytation C10. Slides were first visualized in widefield-brightfield (B,C) or phase contrast mode (not shown) at a 4X magnification. The differing hippocampal regions were noted, and a subregion was selected for further imaging, e.g., dorsal (purple) or ventral (teal). Then, that hippocampal region was imaged at 20X to produce stitched confocal fluorescence images. Slide visualization and annotation enabled the establishment of fiducial markers for navigation and imaging with the CytoViva enhanced-darkfield illumination hyperspectral imaging system. A representative feature from the 20X confocal stitched image is located at 10X on the CytoViva scope. Then, EDF and HSI images are acquired at those regions. This workflow is illustrated in S1A. These images were used to define plaque-associated ROIs for subsequent counting and regional analysis. Scale bars: 2000 μm.

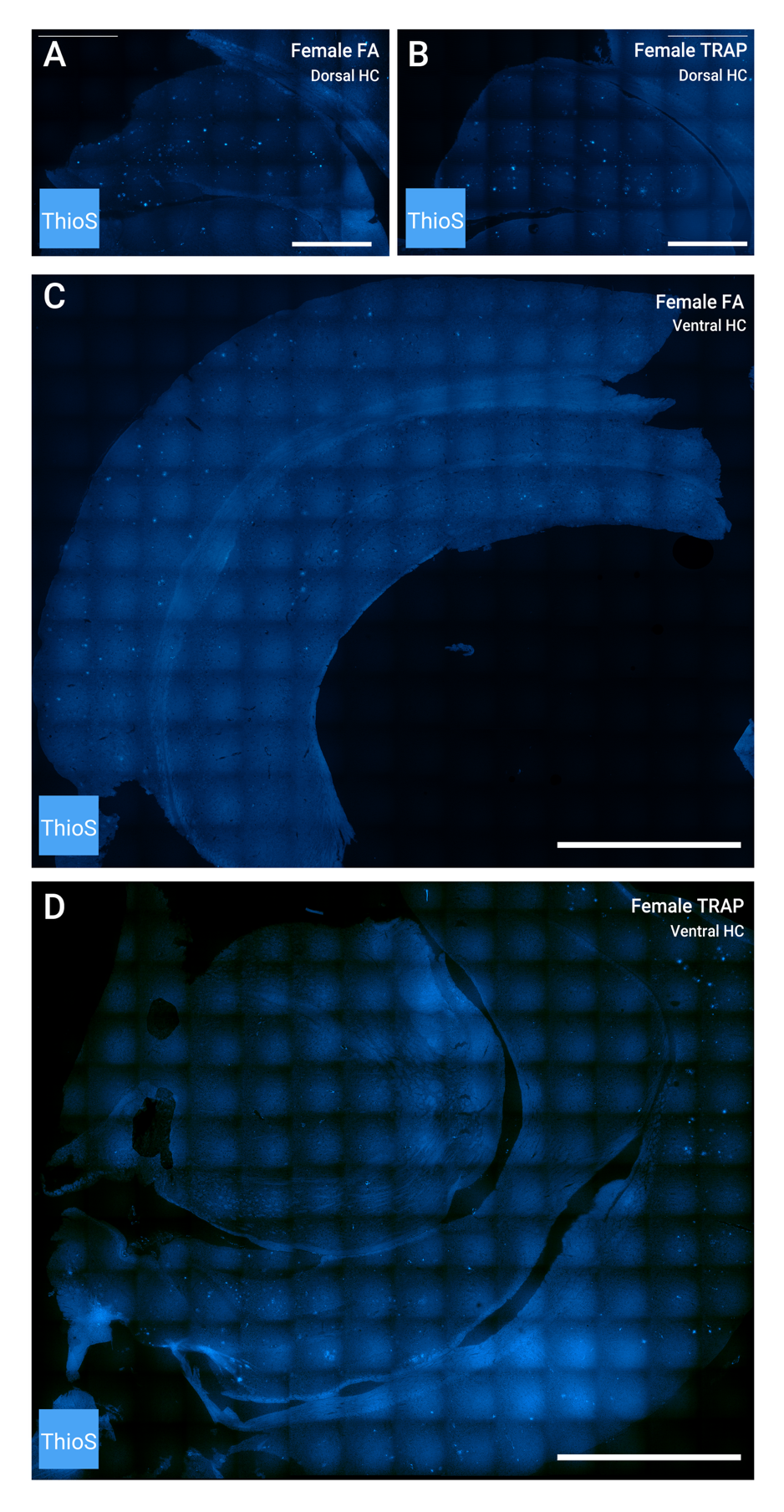

**Supplementary Figure S2. Thioflavin S-stained hippocampal sections used for quantification of Aβ plaque burden in Figure 3.** Representative widefield tiled images of ThioS-stained hippocampal sections from female TgF344-AD rats following 15-month exposure to TRAP or FA. Images include both dorsal (A–B) and ventral (C–D) hippocampal regions, shown to scale. Dorsal hippocampal images from FA-exposed (A) and TRAP-exposed (B) animals correspond to the data shown in Figure 3A–B. Ventral hippocampal sections from matched FA (C) and TRAP (D) animals reveal regional differences in plaque density consistent with the quantifications shown in Figure 3C–E. Such images were used to define plaque-associated ROIs for subsequent counting and regional analysis. Scale bars: 1000 μm.

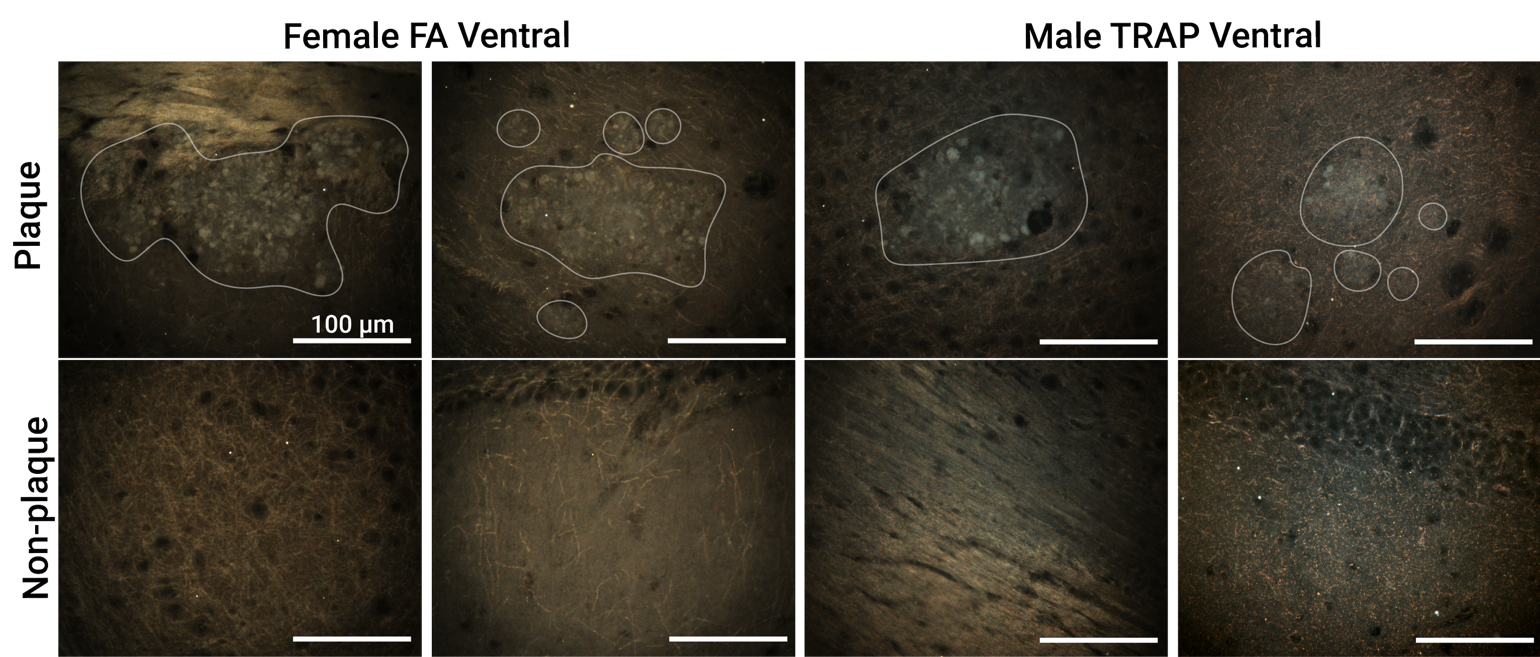

**Supplementary Figure S3. Aβ Plaques visually defer from background tissue in darkfield-enhanced hyperspectral imaging.** Representative EDF-HSI images of four ROIs from the ventral hippocampal region from two TgF344-AD rats (one female, one male), showing the visual distinction of Aβ plaques from background tissue. For further distinction, plaques have been outlined. As explained in the methods section, 8 ROIs in non-plaque regions were chosen per hippocampal region per rat, and 7 ROIs in regions with visible Iba1^+^ fluorescence from confocal imaging were chosen. Those Iba1^+^regions correlated with the presence of Aβ plaques. Particles (bright white puncta) were not contained within Aβ plaque cores, but can be seen proximal to these areas. Scale bars: 100 μm.

**References:**

(1)	Patten, K. T.; Valenzuela, A. E.; Wallis, C.; Berg, E. L.; Silverman, J. L.; Bein, K. J.; Wexler, A. S.; Lein, P. J. The Effects of Chronic Exposure to Ambient Traffic-Related Air Pollution on Alzheimer’s Disease Phenotypes in Wildtype and Genetically Predisposed Male and Female Rats. *Environ Health Perspect* **2021**, *129* (5), 057005. https://doi.org/10.1289/EHP8905.

(2)	Cohen, R. M.; Rezai-Zadeh, K.; Weitz, T. M.; Rentsendorj, A.; Gate, D.; Spivak, I.; Bholat, Y.; Vasilevko, V.; Glabe, C. G.; Breunig, J. J.; Rakic, P.; Davtyan, H.; Agadjanyan, M. G.; Kepe, V.; Barrio, J. R.; Bannykh, S.; Szekely, C. A.; Pechnick, R. N.; Town, T. A Transgenic Alzheimer Rat with Plaques, Tau Pathology, Behavioral Impairment, Oligomeric Aβ, and Frank Neuronal Loss. *J. Neurosci.* **2013**, *33* (15), 6245–6256. https://doi.org/10.1523/JNEUROSCI.3672-12.2013.

(3)	Edwards, S.; Zhao, G.; Tran, J.; Patten, K. T.; Valenzuela, A.; Wallis, C.; Bein, K. J.; Wexler, A. S.; Lein, P. J.; Rao, X. Pathological Cardiopulmonary Evaluation of Rats Chronically Exposed to Traffic-Related Air Pollution. *Environ Health Perspect* **2020**, *128* (12), 127003. https://doi.org/10.1289/EHP7045.

(4)	Patten, K. T.; González, E. A.; Valenzuela, A.; Berg, E.; Wallis, C.; Garbow, J. R.; Silverman, J. L.; Bein, K. J.; Wexler, A. S.; Lein, P. J. Effects of Early Life Exposure to Traffic-Related Air Pollution on Brain Development in Juvenile Sprague-Dawley Rats. *Transl Psychiatry* **2020**, *10* (1), 166. https://doi.org/10.1038/s41398-020-0845-3.

(5)	D. Edelstein, A.; A. Tsuchida, M.; Amodaj, N.; Pinkard, H.; D. Vale, R.; Stuurman, N. Advanced Methods of Microscope Control Using μManager Software. *JBM* **2014**, *1* (2), 1. https://doi.org/10.14440/jbm.2014.36.

(6)	Marlatt, M. W.; Bauer, J.; Aronica, E.; Van Haastert, E. S.; Hoozemans, J. J. M.; Joels, M.; Lucassen, P. J. Proliferation in the Alzheimer Hippocampus Is Due to Microglia, Not Astroglia, and Occurs at Sites of Amyloid Deposition. *Neural Plasticity* **2014**, *2014*, 1–12. https://doi.org/10.1155/2014/693851.

(7)	Schindelin, J.; Arganda-Carreras, I.; Frise, E.; Kaynig, V.; Longair, M.; Pietzsch, T.; Preibisch, S.; Rueden, C.; Saalfeld, S.; Schmid, B.; Tinevez, J.-Y.; White, D. J.; Hartenstein, V.; Eliceiri, K.; Tomancak, P.; Cardona, A. Fiji: An Open-Source Platform for Biological-Image Analysis. *Nat Methods* **2012**, *9* (7), 676–682. https://doi.org/10.1038/nmeth.2019.
